# Notch signaling stabilizes lengths of motile cilia in multiciliated cells in the lung

**DOI:** 10.1101/2024.12.12.628112

**Authors:** Neenu Joy, Aditya Deshpande, Sai Manoz Lingamallu, Vasam Manjveekar Prabantu, CN Naveenkumar, K Bharathkumar, Sukanya Bhat, Zabdiel Alvarado-Martinez, Alessandra Livraghi-Butrico, James S. Hagood, Richard C. Boucher, Daniel Lafkas, Kevin M. Byrd, Shridhar Narayanan, R.K. Shandil, Arjun Guha

**Affiliations:** Institute for Stem Cell Science and Regenerative Medicine (inStem), Bangalore, India – 560065; SASTRA Deemed University, Tirumalaisamudram, Thanjavur, India – 613401; The University of Trans-Disciplinary Health Sciences and Technology (TDU), Yelahanka, Bangalore, India – 560064; Manipal Academy of Higher Education (MAHE), Madhav Nagar, Manipal, India – 576104 54544; Foundation for Neglected Disease Research [FNDR] Plot 20A, KIADB Industrial Area, Doddaballapur, Bangalore-561203; Marsico Lung Institute/Cystic Fibrosis Research Center, University of North Carolina at Chapel Hill, Chapel Hill, North Carolina, USA; Department of Pediatrics (Pulmonology), Marsico Lung Institute, Children’s Research Institute, University of North Carolina at Chapel Hill; Immunology Discovery, Genentech Inc., South San Francisco, CA 94080, USA; Lab of Oral & Craniofacial Innovation (LOCI), Department of Innovation & Technology Research, ADA Science & Research Institute, Gaithersburg, MD, USA; Department of Oral and Craniofacial Molecular Biology, Philips institute for Oral Health Research, Virginia Commonwealth University, Richmond, VA, USA

**Keywords:** Lung, multiciliated cells, ciliary length, Notch signaling, *Mycobacterium tuberculosis* (*M. tb*) infection, host-pathogen interactions

## Abstract

Airway multiciliated cells (MCs) maintain respiratory health by clearing mucus and trapped particles through the beating of motile cilia. While it is known that ciliary lengths decrease along the proximal-distal (P-D) axis of the tracheobronchial tree, how this is regulated is unclear. Here, we demonstrate that canonical Notch signaling in MCs plays a critical role in stabilizing ciliary length. Inhibition of Notch signaling in MCs results in ciliary shortening in the trachea, lengthening in the distal airway, and to region-specific alterations in gene expression. We probe how environmental challenges impact MC homeostasis using germ-free and *Mycobacterium tuberculosis* (*M. tb*) infection models. While germ-free conditions do not perturb ciliary lengths, *M. tb* infection leads to lengthening of distal airway cilia, correlating with a downregulation of Notch signaling. These findings reveal that ciliary length and the P-D gradient in the airways are actively regulated, with Notch signaling serving as a stabilizing mechanism.

## INTRODUCTION

Multiciliated cells (MCs) bear motile cilia that beat in a concerted manner to coordinate fluid flow across epithelial tissues in the lung, brain, and reproductive tract. Motile cilia are hairlike organelles and each cilium is a projection of the plasma membrane comprising a central microtubule scaffold with a ring of 9 doublet protofilaments that surround a central pair of singlet protofilaments (1). This cytoskeletal scaffold, or axoneme as it is called, is tethered to the plasma membrane by a cytoplasmic, centriole-derived basal body located at its base (2). The numbers of motile cilia borne by an MC vary from tens to hundreds, depending on the cell type. Dynein motors located along the ring of microtubule doublets in the ciliary shaft induce relative sliding of these protofilaments to generate a ciliary beat and drive fluid flow (3)

The epithelial lining of the airways is a mucociliary escalator. This escalator comprises MCs and non-ciliated secretory cells. Together, the cells produce airway surface fluid consisting of a cell-proximal periciliary fluid layer and a more luminal, gel-like, mucus layer. The tips of motile cilia contact the mucus layer and, with every beat, displace the mucus layer upwards. The mucus and mucus-trapped particles, including inhaled pathogens, are then swallowed and cleared (4). An integral aspect of the escalator that is conserved across species is the pattern of MCs along the proximal-distal (P-D) axis. MCs in the trachea harbour longer cilia than the MCs in the distal airway (5). This anatomical difference also correlates with differences in the frequencies at which cilia beat (6–8).

The mechanisms that pattern and maintain the mucociliary escalator are not well understood. The specification of MCs during development is orchestrated by a transcriptional hierarchy involving the genes GEMC1, MCIDAS, FOXJ1 and RFX that act in concert with other transcription factors (9–12). Together, these proteins induce a specialized multiciliogenic cell cycle program that coordinates production of ciliary components and assembly of cilia (13). How this program establishes MC heterogeneity along the P-D axis is unclear. A recent study suggests Notch signaling in MCs induces a gradient of Prominin 1 expression along the P-D axis that, in turn, regulates ciliary length (8). The Notch pathway is an evolutionarily conserved, juxtacrine signaling system that plays essential roles in the specification of cell fate (14). The pathway is activated when membrane-bound ligands (Delta-like 1,3,4, Jagged1, 2) bind to membrane-bound Notch receptors (Notch1-4) and trigger cleavage of Notch intracellular domain (NICD) enabling its translocation into the nucleus. Once inside the nucleus, NICD associates with its downstream effector gene *Rbpjk*, and drives the expression of its target genes (14). Prominin 1 (Prom1, CD133) is a cell surface protein with multifarious functions. Pertinently, Prominin 1 may inhibit ciliary growth in a dose-dependent manner (8).

Although developmental programs for MC specification are known, the processes that regulate MC homeostasis post development are only beginning to be unravelled. Exposure of cultures of airway epithelial MCs to cigarette smoke led to a reduction in ciliary length (15). This implies that ciliary length in differentiated MCs is not fixed and can be regulated. The decrease in ciliary length in these cultures is associated with a reduction in the expression of *Foxj1* and *Rfx*, transcription factors that are known to regulate MC development, and in the expression of many ciliary components (16).

We were alerted to a role for the Notch signaling pathway in MC homeostasis during our studies on the role of the pathway in the regulation of non-ciliated secretory cells. Several groups, including ours, have established that Notch signaling is essential for the maintenance of secretory club cells (CCs) in the adult lung and that downregulation of signaling results in the transdifferentiation of the vast majority of CCs into MCs (17–19). Importantly, we noted that Notch2, the predominant Notch receptor expressed in the airway epithelium, is expressed by and activated in virtually all MCs. When we examined airway MCs after Notch inhibition, after transdifferentiation of CCs into MCs, we observed widespread changes in ciliary length and the complete absence of a P-D gradient. These findings led us to probe the role of Notch signaling in MCs and ciliary length.

## RESULTS

### Antibody-mediated inhibition of Notch signaling alters ciliary length and abolishes the P-D gradient and is reversible

The lower airway epithelium of the murine respiratory tract, from trachea to terminal bronchioles (Figure 1A(i)), is composed of several epithelial cell types. MCs and non-ciliated secretory cells (CCs) are the most abundant. MCs are intricately patterned along the P-D axis of the airways with cells in proximal airways possessing more numerous and longer motile cilia (Figure 1A(ii), Figure S1-1). Several groups, including ours, have established that canonical Notch signaling has an indispensable role in airway homeostasis. The pathway is active in CCs and is essential for the maintenance of the CC fate (17–19). Inhibition of Notch signaling results in the transdifferentiation of the vast majority of CCs to MCs (18,19).

**Figure 1:**
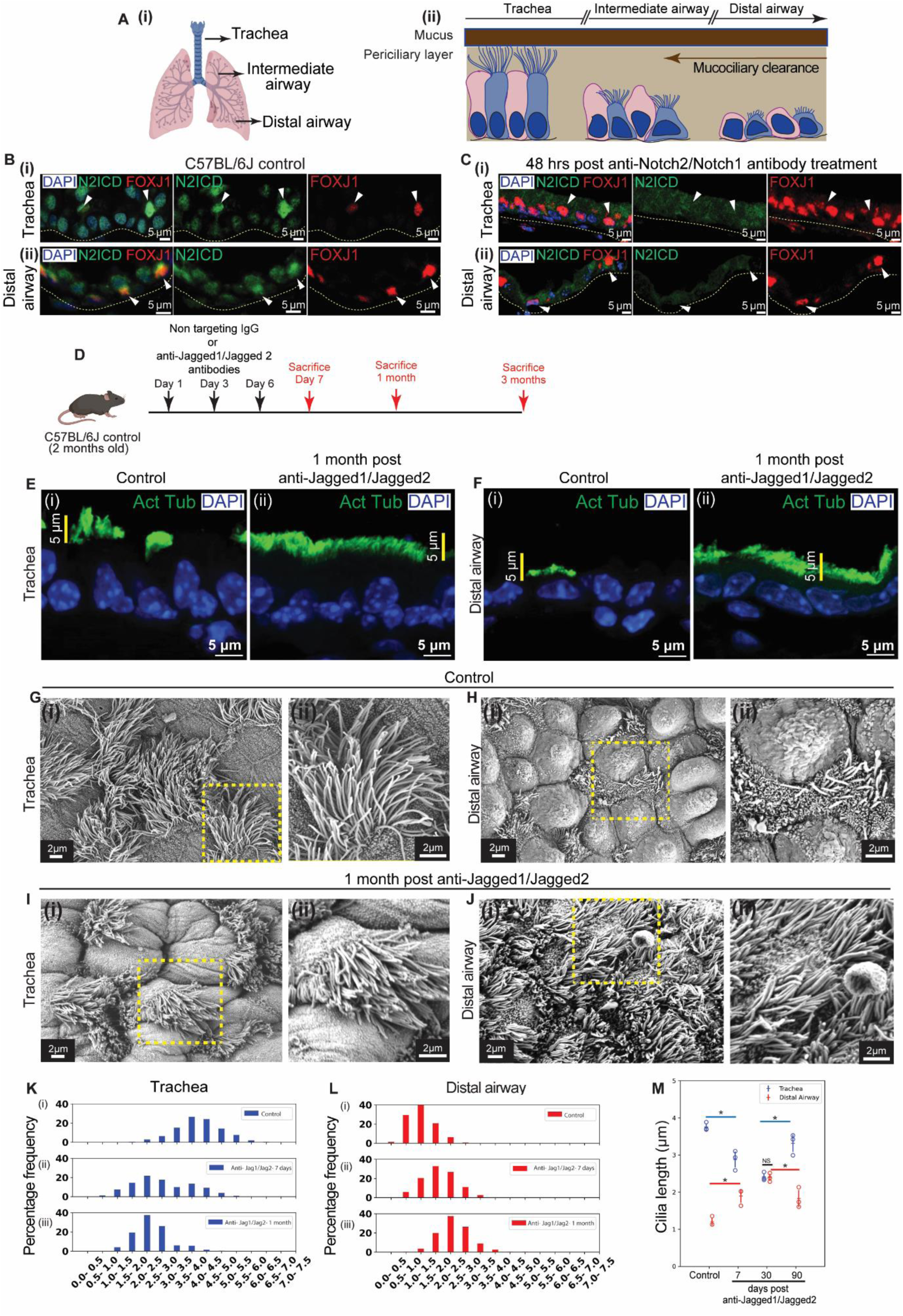
Acute inhibition of Jagged1 and Jagged2 alters ciliary lengths and abolishes the P-D gradient in a reversible manner. **(A)** Diagrams of (i) lung showing trachea, intermediate airways and distal airway and (ii) airway epithelium showing the mucociliary escalator comprising multiciliated cells (MCs, blue) and Club cells (CCs, pink). **(B-C)** Status of Notch signaling in MCs in the adult lung. (B(i)) Tracheal section from C57BL/6J animals showing the distribution of Notch2 intracellular domain (N2ICD, green) in MCs (stained with anti-FOXJ1, red, white arrowheads). Nuclei stained with DAPI (blue). (B(ii)) Lung section from C57BL/6J animals showing the distribution of N2ICD (green) in MCs (stained with anti-FOXJ1, red, white arrowheads) in distal airway. (C(i)) Tracheal section from C57BL/6J animals 48h post anti-Notch1/Notch2 showing N2ICD (green) in MCs (red, white arrowheads). (C(ii)) Lung section from C57BL/6J animals 48h post treatment with anti-Notch1/Notch2 antibodies showing N2ICD (green) in MCs (red, white arrowheads) in distal airway. **(D)** Regimen for inhibition of Notch signaling using non-targeting IgG (control) and anti-Jagged1/Jagged2 antibodies. **(E-F)** Ciliary staining in trachea (E) and distal airway (F) of control (1 month post injection of non-targeting IgG, i) and anti-Jagged1/Jagged2 treated (ii) lungs. (E(i), F(i)) Distribution of anti-Acetylated tubulin (green) in airways from control lungs. (E(ii), F(ii)) Distribution of anti-Acetylated tubulin (green) in airways from anti-Jagged1/Jagged2 treated lungs. Note the pattern of ciliary staining in trachea and distal airway in controls and the absence of this pattern 1 month post antibody treatment. **(G-J)** Scanning electron microscopy of sections from control (1 month post injection of non-targeting IgG) and anti-Jagged1/Jagged2 treated lungs. (G-H) Micrographs of trachea (G(i)-G(ii)) and distal airway (H(i)-H(ii)) from control. Magnified images of boxed regions in (i) are shown in (ii). (I-J) Micrographs of trachea (I(i)-I(ii)) and distal airway (J(i)-J(ii)) from anti-Jagged1/Jagged2 treated lungs. Magnified images of boxed regions in (i) are shown in (ii). **(K-M)** Quantification of ciliary length in micrographs (µm, see methods). **(K-L)** Frequency distribution of ciliary lengths in trachea in control and anti-Jagged1/Jagged2 treated lungs. (K(i)-K(iii)) Frequency distribution of ciliary lengths (µm) in the trachea (blue) in control (i, n=981 cilia), 7 days post anti-Jagged1/ Jagged 2 (ii, n=406 cilia, see Figure S1-2) and 1 month post anti-Jagged1/Jagged2 (iii, n=365 cilia). (L(i)-L(iii)) Frequency distribution of ciliary lengths (µm) in the distal airway (red) in control (i, n=869 cilia), 7 days post anti-Jagged1/ Jagged 2 (ii, n=2409 cilia, see Figure S1-2) and 1 month post anti-Jagged1/Jagged2 (iii, n=954 cilia). Note the alteration of ciliary lengths and absence of a P-D gradient anti-Jagged1/Jagged2 treated lungs. **(M)** Dot plots of ciliary lengths in trachea (blue) and distal airway (red) from control lungs and anti-Jagged 1/Jagged 2 treated lungs 7 days, 1 month and 3 months post treatment (see text for details, n=892 cilia in trachea and n=2430 cilia in distal airway). Hollow circles indicate mean ciliary length in individual animals. Data in dot plot represent mean ± standard deviation (n= 3 animals), (NS-non significant), (* denotes p<0.05, Student’s t-test). Note that the alteration of ciliary lengths post anti-Jagged1/Jagged2 treatment is reversed by 3 months. See also Figure S1-1 to Figure S1-3.

The status of canonical Notch signaling in airways has been assessed by staining lung sections with an antibody that detects the intracellular domain of Notch2 (N2ICD), the predominant Notch receptor expressed in the airways (18). N2ICD translocates to the nucleus upon activation and cells that activate Notch signaling exhibit nuclear localized N2ICD. Lung sections from adult mice (2-3 months of age) stained with anti-N2ICD revealed that the protein is nuclear localized in CCs and, unexpectedly, also in MCs (Figure 1B (i-ii)). To test the specificity of N2ICD staining in MCs, we inhibited Notch signaling by injecting mice anti-Notch2/Notch1 or anti-Jagged1/Jagged2 antibodies (19), isolated lungs 48 hours later and stained lung sections with anti-N2ICD (Figure 1C). Under these conditions we observed a complete loss of nuclear N2ICD in both CCs and MCs (Figure 1C (i-ii)), showing that N2ICD immunostaining in MCs was specific. Next, we stained lung sections from older mice with anti-N2ICD (aged 1.2-1.5 years). Nuclear N2ICD was detected in virtually all MCs (n=3 animals, data not shown). We inferred that Notch signaling is activated in MCs in the adult lung.

The role of canonical Notch signaling in differentiated airway MCs has not been characterized. Thus, we decided to investigate whether Notch signaling has any role in MCs. Mice were injected with either non-targeting Immunoglobulin G (IgG control) or anti-Jagged1/Jagged2 antibodies (for detailed protocol see methods) and tracheae and lungs from these animals were harvested at different time points and processed for histological analysis (schematic in Figure 1D). The structure of cilia on MCs was analysed by fluorescence and scanning electron microscopy (SEM).

Immunostaining with anti-Acetylated tubulin, an antibody that detects post-translationally modified tubulin enriched in motile cilia, clearly outlines MCs in lung sections (Figure S1-1, Figure 1E(i)-F(i)). The antibody illuminates a band of the luminal cilia, diminishing in thickness along the P-D axis (Figure S1-1, Figure 1E(i)-F(ii)). Sections from lungs treated with either non-targeting IgG or anti-Jagged1/Jagged2 antibodies were stained with anti-Acetylated tubulin and examined. We detected a lawn of Acetylated tubulin staining in sections from anti-Jagged treated lungs at both 7 days and 1 month after injection (Figure 1E(ii),F(ii), (18,19)). Interestingly, differences in the thickness of luminal staining along the P-D axis were perturbed at 7 days post treatment (not shown) and conspicuously absent 1 month after antibody treatment (Figure 1E(ii),F(ii)). These stainings suggested that the inhibition of Notch signaling had altered ciliary length in all MCs throughout the airways. Proximal cilia appeared to decrease in length while distal airway cilia increased in length (compare Figure 1E(i)–1E(ii) and 1F(i)-F(ii)).

In order to examine the effects of Notch inhibition at higher resolution we turned to SEM. Thick sections (200μm) of tracheal rings and lung (left lobe) were imaged. Motile cilia were observed throughout the airways and ciliary length appeared to decrease along the P-D axis (Figure S1-1). Ciliary lengths in the trachea and distal airways (terminal bronchioles) were quantified from micrographs using a free-hand contour-tracing approach (see methods). Measurements of length demonstrated a statistically significant decrease in ciliary length from trachea through intermediate airways to terminal bronchioles (Figure S1-1G). Next, we compared ciliary morphology in lungs from mice that were injected with either non-targeting IgG or anti-Jagged1/Jagged2 antibodies (Figure 1G-M). Anti-Jagged treatment led to a decrease in ciliary length in the trachea and to an increase in ciliary length in the distal airway (Figure 1G-J, histograms of ciliary lengths shown in Figure 1K-L, representative images from 7 days shown in Figure S1-2, dot plots of ciliary lengths in trachea and distal airway across all time points shown in Figure 1M). No P-D gradient in ciliary length was detected at 1 month (Figure 1M).

Previous studies have shown that the perturbation in the balance of cell types induced by anti-Jagged treatment is gradually reversed. Over time Notch signaling is restored and variant club cells, rare CCs that resist transdifferentiation into MCs, proliferate and repopulate the airways with CCs and MCs (19). At 3 months post treatment, patches of cells comprising CCs and MCs could be detected in the lawn of MCs (19). In order to probe the fate of MCs and cilia post anti-Jagged treatment long-term, we utilized a lineage tracing approach. For this, *FoxJ1*^CreERT2/+^ mice, that drive expression of the creER-recombinase specifically in MCs, were bred to Rosa^Tdtomato/+^ animals. The *FoxJ1*^CreERT2/+^; Rosa^Tdtomato/+^ progeny were injected with anti-Jagged1/Jagged2 antibodies to induce transdifferentiation and then injected with tamoxifen to induce Tdtomato (Tdt) reporter expression in both pre-existing and transdifferentiated MCs (hereafter collectively referred to as “old MCs”, experimental pipeline in Figure S1-3A). The lineage-labeled animals were then sacrificed 3 months post treatment and the distribution of MCs was examined. Old MCs (Tdt+) were detected in a lawn throughout the airways and dispersed within this lawn were patches of Tdt- cells comprising CCs and new MCs (Figure S1-3B-E). We inferred that the old MCs persisted till 3 months and were the predominant population in the airways at this time point. Next, we imaged MCs in these lungs using SEM taking care to avoid patches of cells comprising both CCs and MCs (Figure S1-3F,G). Quantitative analysis of ciliary lengths 3 months post anti-Jagged treatment showed that, in comparison to the lengths at 1 month, tracheal ciliary length had increased and distal airway ciliary length had decreased (Figure 1M).

Taken together, these findings led us to the following conclusions. First, MCs activate Notch signaling during homeostasis. Second, the signaling pathway regulates ciliary length throughout the airways and inhibition of the pathway abolishes the P-D gradient in ciliary length. Third, changes in ciliary length that occur upon acute Notch inhibition are reversible once signaling is restored.

### Genetic ablation of *Rbpjk* in multiciliated cells (MCs) establishes a role for canonical Notch signaling in maintaining ciliary length during homeostasis

To investigate whether Notch signaling in MCs specifically regulates ciliary homeostasis, we adopted a genetic approach. We conditionally deleted *Rbpjk*, the transcription factor that orchestrates changes in gene expression in the canonical Notch signaling pathway, in MCs and examined the impact on MC morphology. For this, *FoxJ1*^CreERT2/+^ mice were bred to *RBPJκ*^flox/flox^, *FoxJ1*^CreERT2/+^; *RBPJκ*^flox/flox^ were generated and induced with Tamoxifen to delete *Rbpjk* (hereafter called as *Rbpjk*-flox/flox). Littermates that did not receive tamoxifen we utilized as controls (experimental pipeline shown in Figure 2A). The efficiency of *Rbpjk* deletion in *Rbpjk*-flox/flox animals was determined by immunostaining lung sections for RBPJk at 10 days and 1 month post tamoxifen administration (Figure S2-1). Nuclear expression of Rbpjk was observed in all the airway epithelial cells in controls, including MCs (Figure S2-1B, S2-1C, S2-1D), and depleted in *Rbpjk*-flox/flox at both 10 days and 1 month (Figure S2-1B, S2-1E, S2-1F). We noted that the efficiency of *Rbpjk* depletion in MCs was lower in the trachea than in the distal airway (terminal bronchioles) and higher overall at 1 month than 10 days post induction (Figure S2-1B).

**Figure 2:**
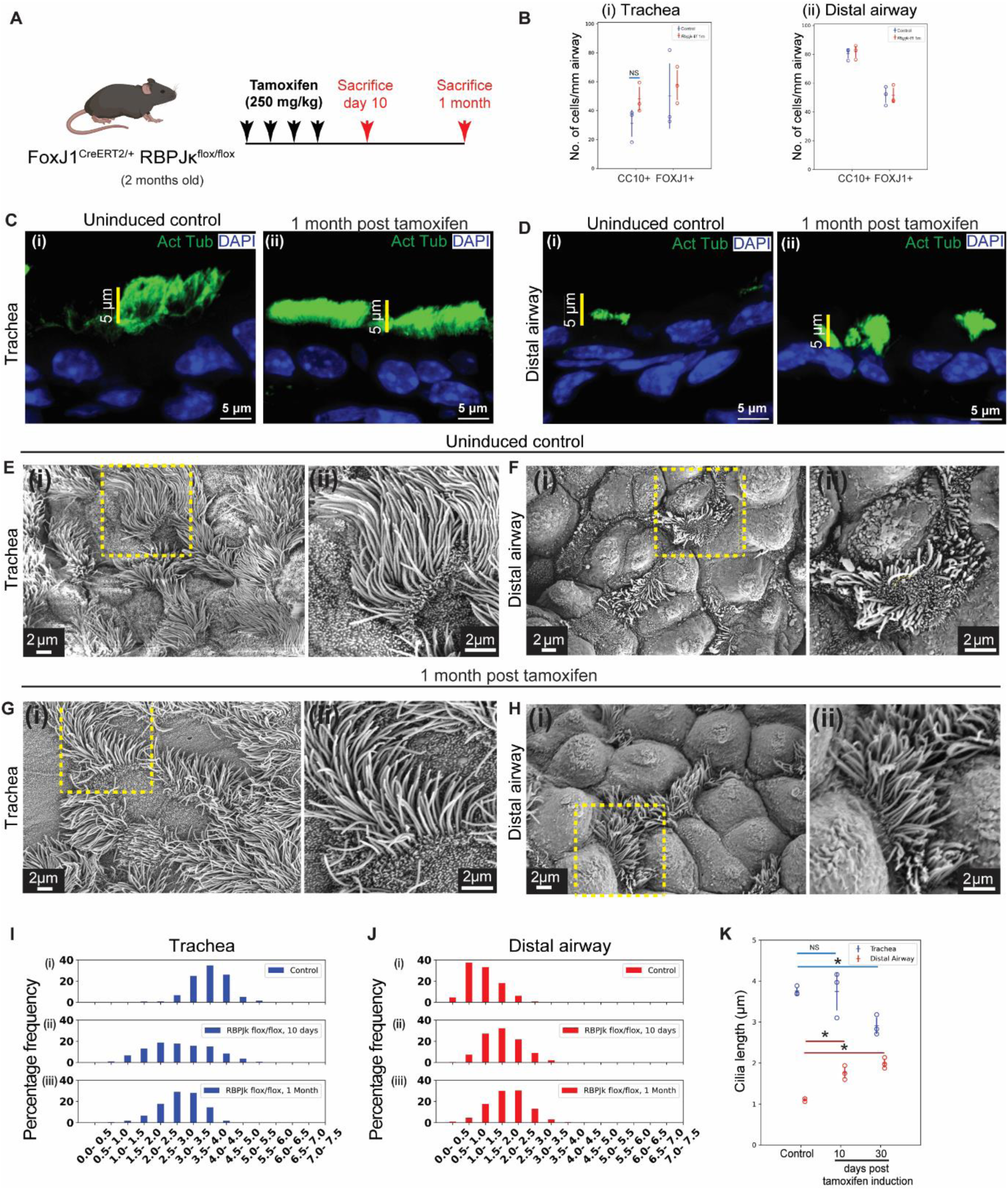
Genetic ablation of *Rbpjk* in MCs alters ciliary lengths and the P-D gradient. **(A)** Outline of the genetic strategy used to conditionally delete *RBPJκ* in MCs. (**B)** Dot plots showing frequencies of club cells (CC10+) and MCs (FOXJ1+) in trachea (B(i)) and distal airway (B(ii) from control (uninduced, blue) and mutant (induced, “*Rbpjk*-flox/flox”, red) lungs. Hollow circles indicate mean cell frequency/mm airway in immunostained sections from an individual animal (see text, methods). Data in dot plot represent mean ± standard deviation (n=3 animals) **(C-D)** Ciliary staining in trachea (C) and distal airway (D) of control (age matched uninduced littermate, (i)) and *Rbpjk*-flox/flox (ii) lungs. (C(i),D(i)) Distribution of anti-Acetylated tubulin (green) in airways from control lungs. (C(ii),D(ii)) Distribution of anti-Acetylated tubulin (green) in airways from *Rbpjk*-flox/flox lungs. Note the pattern of ciliary staining in trachea and distal airway in control and the absence of this pattern in *Rbpjk*-flox/flox 1 month post induction. **(E-H)** Scanning electron microscopy of sections from control (age matched uninduced littermate) and *Rbpjk*-flox/flox lungs. **(E-F)** Micrographs of trachea (E(i)-E(ii)) and distal airway (F(i)-F(ii)) from control lungs. Magnified images of boxed regions in (i) are shown in (ii). **(G-H)** Micrographs of trachea (G(i)-G(ii)) and distal airway (H(i)-H(ii)) from *Rbpjk*-flox/flox lungs. Magnified images of boxed regions in (i) are shown in (ii). **(I-K)** Quantification of ciliary length in micrographs (µm, see methods). **(I-J)** Frequency distribution of ciliary lengths in trachea in control and *Rbpjk*-flox/flox lungs. (I(i)-I(iii)) Frequency distribution of ciliary lengths (µm) in the trachea (blue) in control (age matched uninduced littermate, i, n=667 cilia), 10 days post induction (ii, n=535 cilia) and 1 month post induction (iii, n=685 cilia). (J(i)-J(iii)) Frequency distribution of ciliary lengths (µm) in the distal airway (red) in control (age matched uninduced littermate, i, n=1795 cilia), 10 days post induction (ii, n=733 cilia) and 1 month post induction (iii, n=1173 cilia). Note the alteration of ciliary lengths and reduction of the P-D gradient in *Rbpjk*-flox/flox lungs. **(K)** Dot plots of ciliary lengths in trachea (blue) and distal airway (red) in control (age matched uninduced littermate), *Rbpjk*-flox/flox 10 days and 1 month post tamoxifen induction. Hollow circles indicate mean ciliary length in individual animals. Data in dot plot represent mean ± standard deviation (n= 3 animals), (NS-non significant), (* denotes p<0.05, Student’s t-test). See also Figure S2-1 to Figure S2-2.

Having established the extent of loss of *Rbpjk* in *Rbpjk*-flox/flox, we proceeded to characterize the impact of *Rbpjk* depletion on the overall frequencies of MCs and CCs in the airway epithelium. For this, we performed immunostainings for CC10/Scgb1a1 (a marker for CCs), and FoxJ1 (MCs) and quantified the frequencies of these cells in the trachea and distal airway at 1 month in controls and *Rbpjk*-flox/flox. The frequencies of these cell types in trachea and the distal airway were comparable (Figure 2B). We infer that loss of Notch signaling in MCs does not alter the balance of MCs and CCs in the airway epithelium.

Next, we probed the effect of *Rbpjk* ablation on the morphology of MCs. For this, we performed immunostaining for Acetylated-tubulin and examined thick sections using SEM. Immunostaining revealed a qualitative decrease in ciliary length in trachea (compare Figure 2C(i)– 2C(ii)) and an increase in length in distal airway (compare Figure 2D(i)- 2D(ii)) in *Rbpjk*-flox/flox at 1 month. No consistent difference in ciliary length was observed at 10 days (data not shown). SEM imaging of control and *Rbpjk*-flox/flox and quantitation of ciliary length across time points showed that ciliary length decreased in the trachea and increased in the distal airway (Figure 2 E-K, histograms of ciliary lengths shown in Figure 2I-J, representative images from 10 days shown in Figure S2-2, dot plots of ciliary lengths in trachea and distal airway across all time points shown in Figure 2K). We infer that canonical Notch signaling in MCs stabilizes ciliary length and the P-D gradient. The extent of the perturbation post anti-Jagged1/Jagged2 treatment (Figure 1) is greater than in *Rbpjk*-flox/flox (Figure 2). The difference could be due to the incomplete penetrance of *Rbpjk* deletion in the *Rbpjk*-flox/flox (Figure S2-1).

### Spatial transcriptomics suggests that Notch signaling in MCs has a pervasive role in regulating airway gene expression and homeostasis

Having established that canonical Notch signaling in MCs has a role in the regulation of MC morphology, next we examined how this axis of Notch signaling impacts airway homeostasis. We adopted a spatial transcriptomic approach using the GeoMX Digital spatial profiler (DSP) platform that facilitates analysis of gene expression in specific regions of interest (ROIs). Paraffin sections of trachea and lung lobes from control (*FoxJ1*^CreERT2/+^; Rosa^Tdtomato/+^ induced with tamoxifen) and *Rbpjk*-flox/flox at 1 month post tamoxifen were hybridized with an RNA probe set, immunostained with anti-Acetylated tubulin antibody to illuminate airways/MCs, and ROIs were extracted from trachea and distal-most airways (6 ROIs from control trachea, 14 ROIs from control distal airway, 12 ROIs from *Rbpjk*-flox/flox mice trachea, and 12 ROIs from *Rbpjk*-flox/flox mice distal airway). Following tissue extraction, probes were isolated from different ROIs for library preparation and sequencing (experimental pipeline shown in Figure 3A, see methods for a detailed protocol).

**Figure 3:**
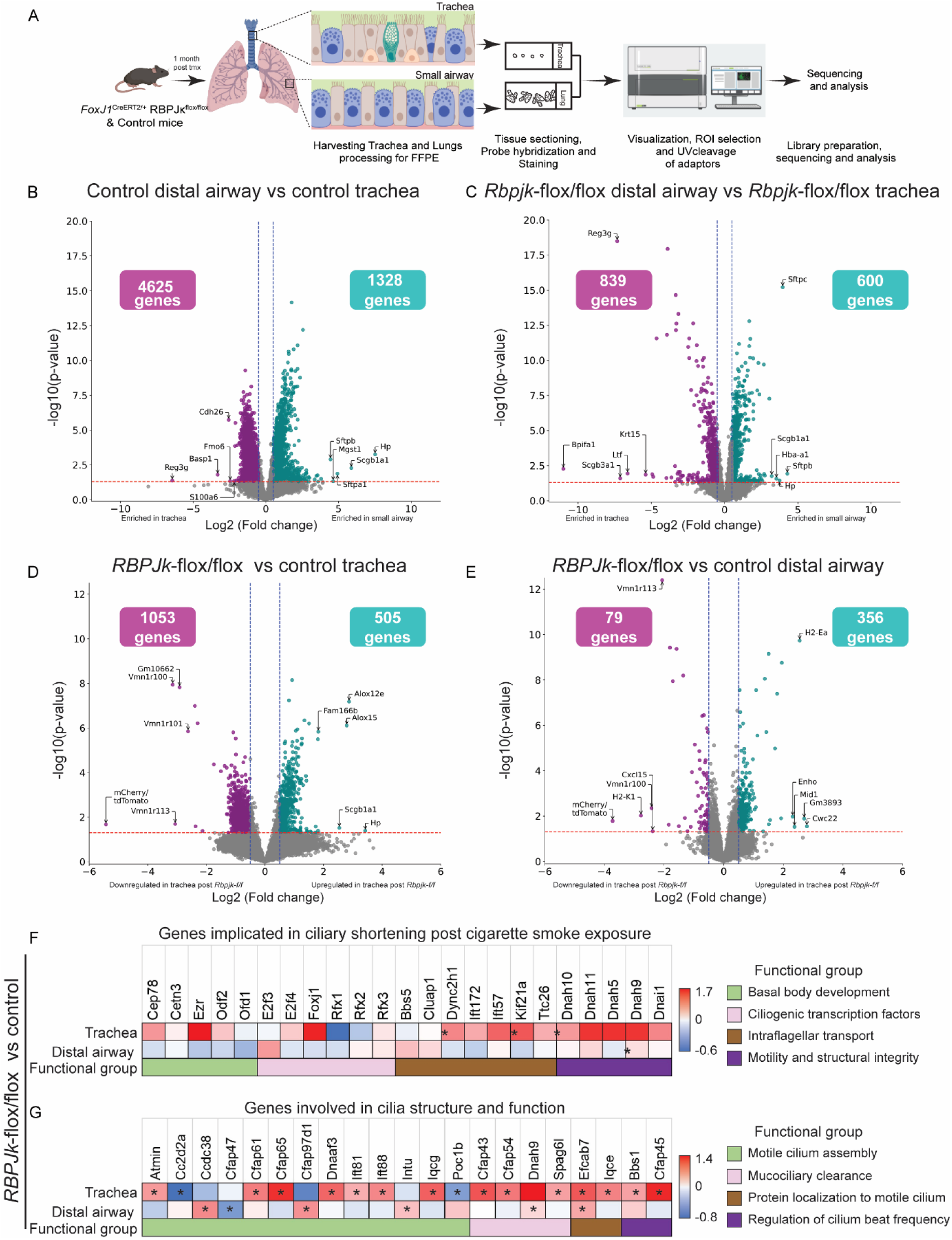
Spatial transcriptomics reveals that Notch signaling in MCs regulates airway gene expression and homeostasis. **(A)** Experimental design for spatial transcriptomics analysis of trachea and distal airway of control and *Rbpjk*-flox/flox mice (also see methods) **(B)** Volcano plot showing differentially expressed genes (DEGs) in trachea and distal airway ROIs in lung sections from control *FoxJ1*^CreERT2/+^; Rosa^Tdtomato/+^ animals. Individual genes (dots) with log2 fold change <-0.5, p<0.05 (purple, enriched in trachea) and log2 fold change>0.5, p<0.05 (cyan, enriched in distal airway) are highlighted. Top 5 enriched genes in trachea and distal airway are labelled. **(C)** Volcano plot showing DEGs in trachea and distal airway ROIs in *Rbpjk*-flox/flox lungs. Individual genes (dots) with log2 fold change <-0.5, p<0.05 (purple, enriched in trachea) and log2 fold change >0.5, p<0.05 (cyan, enriched in distal airway) are highlighted. Top 5 enriched genes in trachea and distal airway are labelled. Note the reduction in frequencies of DEGs in trachea and distal airway in *Rbpjk*-flox/flox lungs. **(D)** Volcano plot showing DEGs in tracheal ROIs from control and *Rbpjk*-flox/flox lungs. Individual genes (dots) with log2 fold change <-0.5, p<0.05 (enriched in control, purple) and log2 fold change >0.5, p<0.05 (enriched in *Rbpjk*-flox/flox, cyan) are highlighted. **(E)** Volcano plot showing DEGs in distal airway ROIs from control and *Rbpjk*-flox/flox lungs. Individual genes (dots) with log2 fold change >0.5, p<0.05 (enriched in *Rbpjk*-flox/flox, cyan) or log2 fold change <-0.5, p<0.05 (enriched in control, purple) are highlighted. The number of highlighted genes in control and *Rbpjk*-flox/flox are given in boxes in the respective volcano plots. (D-E) Top 5 upregulated and downregulated genes are labelled in the respective plots. **(F)** Heat map of genes implicated in ciliary shortening post cigarette smoke exposure in trachea and distal airway of *Rbpjk-*flox/flox mice compared to control. **(G)** Heat map showing genes involved in cilia structure and function (MSigDB REACTOME database, Gene Ontology Biological Processes (GO_BP): terms related to cilia structure and function) in trachea and distal airway of *Rbpjk-* flox/flox mice compared to control. Significantly expressed genes in either trachea or distal airway are plotted. Asterisk (*) denotes p<0.05.

A comparative analysis of gene expression profiles from trachea and distal airway ROIs in control samples revealed the following. First, high levels of *Tdt* were detected in ROIs from controls (Source Data Table 2). Second, the gene expression patterns in tracheal and distal airway ROIs were significantly different (Figure 3B, genes with |log 2-fold change| > 0.5 and p < 0.01 are used). We have demonstrated previously that Reg3g expression is higher in tracheal CCs while Scgb1a1 and Hp expression are higher in distal airway CCs (ref). Gene expression profiles in tracheal and distal airway ROIs recapitulates these differences (Figure 3B, see annotated genes) suggesting that the transcriptomic profiles are indeed representative. Next, we analysed gene expression in tracheal and distal airway ROIs derived from control and *Rbpjk*-flox/flox animals. No Tdt expression was detected in sections from *Rbpjk*-flox/flox, consistent with the absence of the Tdt reporter in this genetic background (Source Data Table 2). We detected differences in gene expression between tracheal and distal airway ROIs. However, there is a difference in the frequencies of differentially expressed genes in *Rbpjk*-flox/flox (Figure 3C). While 5953 genes are differentially expressed across distal airway and tracheal ROIs in controls (Figure 3B), only 1439 genes are differentially expressed in *Rbpjk*-flox/flox (Figure 3C). This suggests that the ablation of Notch signaling in MCs leads to a reduction in airway gene expression.

Next, we compared the differences in gene expression between control and *Rbpjk*-flox/flox in the tracheal and distal airway ROIs respectively (Figure 3D-E). In tracheal ROIs, 505 genes were expressed at higher levels and 1053 genes were downregulated (Figure 3D). In distal airway ROIs, 356 genes were expressed at higher levels and 79 genes were downregulated (Figure 3E). There was little overlap between the lists of genes up or downregulated in trachea and distal airway (list of genes in Source Data Table 1). A closer examination of the lists of differentially expressed genes in *Rbpjk*-flox/flox and control in different airway regions revealed that genes that showed greatest change in levels of expression in either trachea or distal airway typically expressed in cell types other than MCs (Figure 3D-E). For example, among the 5 genes that show highest upregulation in the trachea in *Rbpjk*-flox/flox are *Hp* and *Scgb1a1/CC10*, genes that are typically expressed in CCs (Figure 3D-E). We cross-referenced the lists of differentially expressed genes in trachea and distal airway with lung single cell RNA sequencing data from various sources (20) (AG, NJ, SML, AD, VMP, unpublished) to identify which cells typically express these genes (Figure 3D-E, Source Data Table 1). This analysis shows that many of the differentially expressed genes are expressed in other cells, not in MCs (Source Data Table 1). The transcriptomic analysis shows that Notch signaling in MCs is likely to have a pervasive role in the regulation of airway gene expression and homeostasis.

In light of the role of Notch signaling in stabilizing ciliary length (previous sections), next we examined how perturbations to Notch signaling impact expression of ciliary genes. Exposure to cigarette smoke has been shown to impact MC homeostasis and ciliary length (15, 16). This study correlated the decrease in ciliary length in tracheal MCs with a decrease in the expression of transcription factors that orchestrate ciliogenesis and their target genes (16). To probe the role of Notch signaling in the regulation of ciliary genes, we examined 24 genes shown to be downregulated upon exposure to cigarette smoke (Figure 3F). We do not detect a significant downregulation of any of these genes in the tracheal ROIs (where there is a decrease in ciliary length) nor an upregulation in the distal airway ROIs (where there is an increase in ciliary length) (Source Data Table 3). This suggests that cigarette smoke exposure and perturbations in Notch signaling alter ciliary lengths by different mechanisms. We subsequently examined how perturbations in Notch signaling impact expression of 107 genes associated with ciliogenesis and ciliary function and found that 22 genes showed changes in levels that are statistically significant (Figure 3G, Source Data Table 3, p<0.05). Although we do detect differences in the expression of several ciliary genes in tracheal and distal airway ROIs (Figure 3G), consistent with changes in ciliary morphology, these changes in gene expression are not concerted across airway regions. How Notch-dependent regulation of expression of ciliary genes regulates ciliary length will require investigation (also see Discussion).

### *M. tuberculosis* infection induces multiciliated cell remodelling in distal airway akin to Notch inhibition

Having established a role for Notch signaling in MCs in ciliary and airway homeostasis, next we probed how this regulatory mechanism is impacted by environmental challenges. As mentioned previously, the function of the mucociliary escalator is clearance of mucus and mucus-trapped particles, including pathogens. The escalator is also essential for clearance of excess mucus that results from infection and inflammation. Thus, we decided to probe how the lung microbiome impacts Notch signaling-dependent control of ciliary and airway homeostasis using two approaches. First, we analysed lungs from germ-free mice that lack an intrinsic microbiome. Second, we analysed lungs from mice infected with *Mycobacterium tuberculosis* (*M. tb,* H37Rv strain).

We did not observe any differences in the gradation of ciliary length across the airways of germ-free mice (Figure S4-1). This suggests that the intrinsic airway microbiome does not influence MC development or MC homeostasis. The mouse model of *M. tb* infection is characterized by severe infection, inflammation and formation of granulomas in the parenchyma. In response to aerosolized H37Rv exposure, colony forming unit (CFU) counts in lung peak by 1 month post infection and remains stable thereafter (20). Granulomas consisting of loose, non-necrotic cellular aggregates, with a discrete fibrotic reaction, lack of encapsulation, and strong lymphocyte presence, develop as early as 2-4 weeks after infection (21). We infected female BALB/c mice with *M. tb* and isolated lungs 1 day, 1 month, 3 months, and 6 months post infection to estimate bacterial titres, examine MC morphology and analyse airway gene expression (experimental pipeline shown in Figure 4A, see methods).

**Figure 4:**
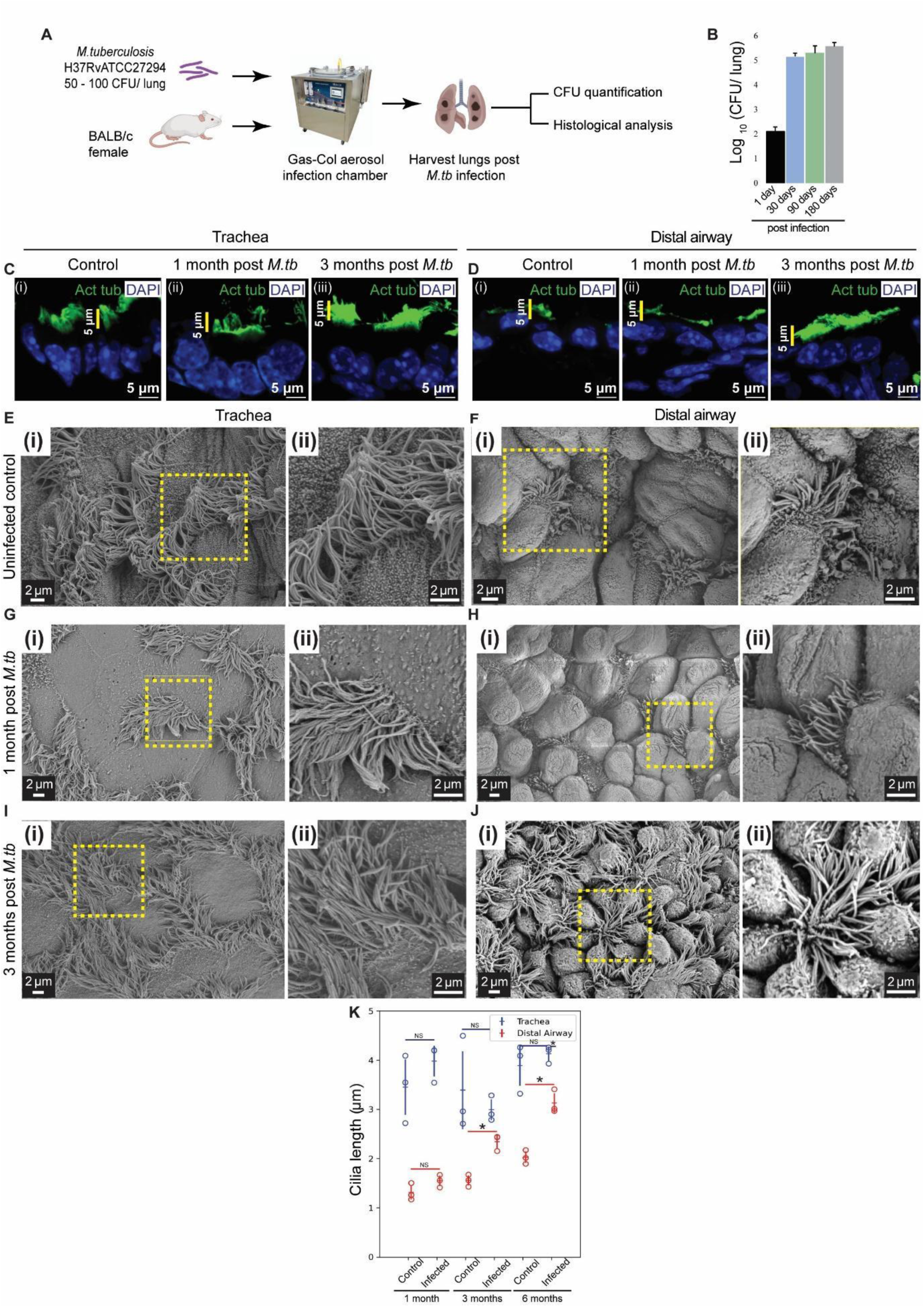
*M. tuberculosis* infection induces multiciliated cell remodelling akin to Notch inhibition in distal airway. **(A)** Experimental design for *M. tuberculosis* (*M. tb*) infection in BALB/c mice **(B)** Frequencies of colony forming units (CFU) of *M. tb* in infected lungs at 1 day, 1 month, 3 months and 6 months post infection (n=3 animals for each time point) **(C-D)** Ciliary staining in trachea (C) and distal airway (D) of control (BALB/c uninfected age matched cohort, i) and *M. tb* infected lungs (ii, iii). (C(i), D(i)) Distribution of anti-Acetylated tubulin (green) in airways from control lungs. (C(ii), D(ii)) Distribution of anti-Acetylated tubulin (green) in airways from *M. tb* 1 month post infection. (C(iii), D(iii)) Distribution of anti-Acetylated tubulin (green) in airways from *M. tb* 3 months post infection. Note the pattern of ciliary staining in trachea and distal airway in controls and the absence of this pattern 3 months post *M. tb* infection. **(E-J)** Scanning electron microscopy of sections from control and *M. tb* infected lungs. **(E-F)** Micrographs of trachea (E(i)-E(ii)) and distal airway (F(i)-F(ii)) from control lungs. Magnified images of boxed regions in (i) are shown in (ii). **(G-H)** Micrographs of trachea (G(i)-G(ii)) and distal airway (H(i)-H(ii)) from lungs 1 month post *M. tb* infection. Magnified images of boxed regions in (i) are shown in (ii). **(I-J)** Micrographs of trachea (I(i)-I(ii)) and distal airway (J(i)-J(ii)) from lungs 3 months post *M. tb* infection. Magnified images of boxed regions in (i) are shown in (ii). **(K)** Dot plot of ciliary lengths in trachea (blue) and distal airway (red) from control lungs (uninfected, age-matched, n=314 in trachea, n=397 in distal airway) and from lungs 1 month (n=353 in trachea, n=645 in distal airway), 3 months (n=356 in trachea, n=1082 in distal airway), 6 months (n=230 in trachea, n=1079 in distal airway, see Figure S4-2) post *M. tb* infection. Hollow circles indicate mean ciliary length in individual animals. Data in dot plot represent mean ± standard deviation (n= 3 animals), (NS-non significant), (* denotes p<0.05, Student’s t-test). See also Figure S4-1 to Figure S4-2.

Estimates of bacterial titres in mice from various time points showed that infection peaked at 1 month and remained stable thereafter (Figure 4B). To assess the impact of pathogen challenge on MC morphology, we once again utilized fluorescence microscopy and SEM. Sections from control and infected mice were immunostained with anti-Acetylated tubulin. We did not detect any changes in the ciliary staining in tracheal sections at 1, 3 and 6 months post infection (compare Figure 4C (i), (ii), (iii), see Figure S4-2A for 6 months post *M. tb* infection). Interestingly, we noted a striking increase in ciliary length, akin to Notch mutants, in the distal airway by 3 months post infection (compare Figure 4D (i), (ii), (iii), see Figure S4-2B for 6 months post *M. tb* infection). Quantification of ciliary lengths in micrographs (SEM) showed that there were no significant differences in tracheal ciliary length at any time points post infection (compare Figures 4E(i), 4G(i), 4I(i), quantified in Figure 4K, see Figure S4-2 for 6 months post *M. tb* infection). Quantification of ciliary length of distal airway showed that there was an increase by 3 months post infection (compare Figures 4F(i), 4H(i), 4J(i), quantified in Figure 4K). Interestingly, the increase in ciliary length in the distal airway was observed in virtually every distal airway examined and did not correlate with proximity to sporadic granulomas. We infer that the alterations in MC and ciliary homeostasis observed in Notch signaling-deficient lungs can also be observed in response to *M*. *tb* infection.

### Alterations in MC homeostasis in *M. tb* infected lungs correlated with downregulation of Notch signaling

The changes in ciliary architecture in the distal airway post Notch inhibition and post *M. tb* infection were similar. This finding led us to probe if Notch signaling is perturbed in the distal airway upon *M. tb* infection. In order to examine the status of Notch signaling in MCs, we immunostained sections from uninfected control and *M. tb*-infected lungs at all time points with anti-N2ICD. MCs in uninfected animals, in both tracheal and distal airway, contained nuclear N2ICD at all time points. MCs in *M. tb* infected animals showed a different pattern (Figure 5A-F). Although MCs in the trachea contained nuclear N2ICD at all time points examined, MCs in the distal airway did not. N2ICD staining was markedly downregulated in distal airway by 3 months (Figure 5F). This suggests that Notch signaling is downregulated in distal airway MCs 3 months post *M. tb* infection.

**Figure 5:**
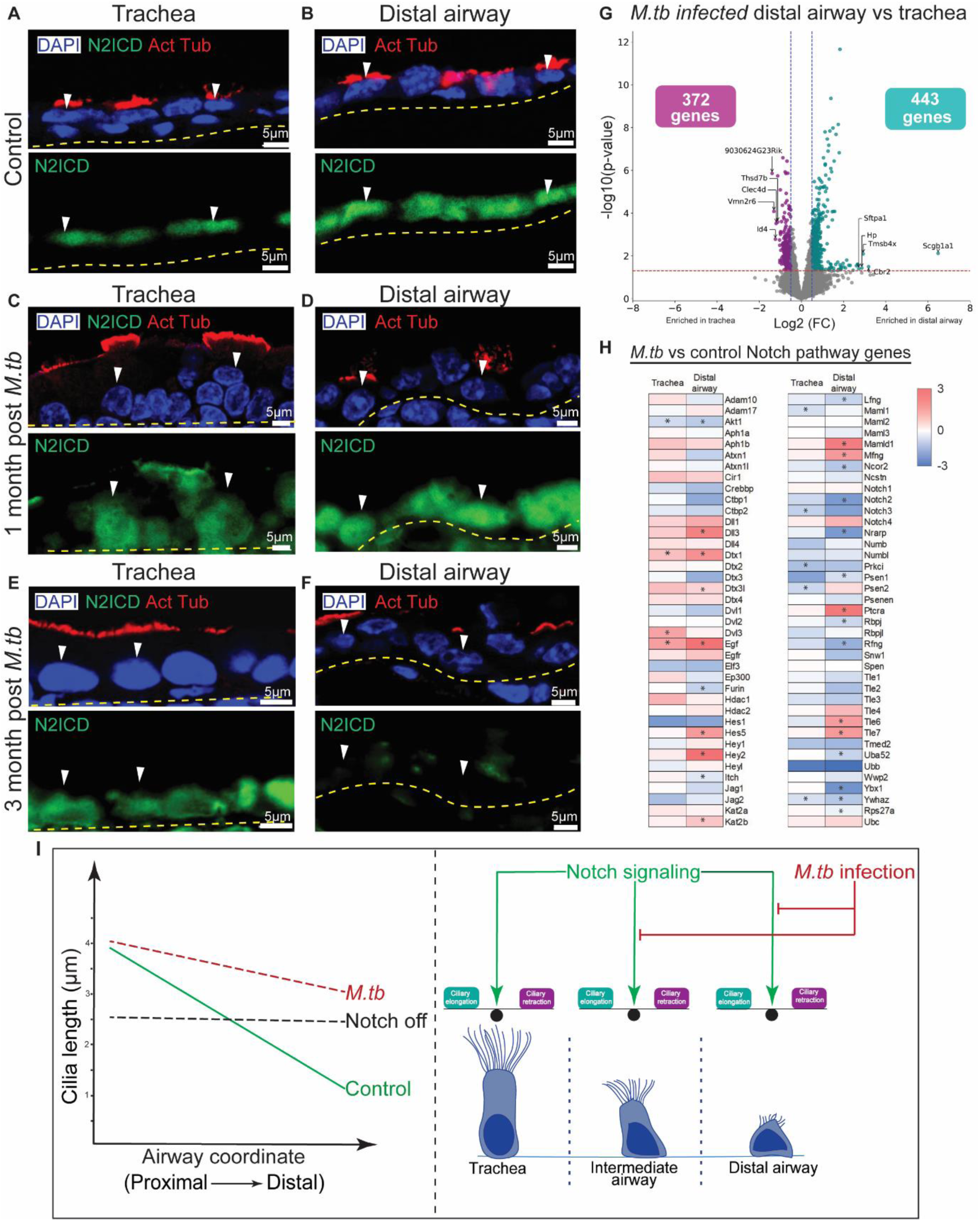
*M. tuberculosis* infection perturbs Notch signaling in distal airway. **(A-F)** Status of Notch signaling in MCs in uninfected and *M. tb* infected lungs. (A,C,E) Tracheal sections from control (BALB/c age matched uninfected cohort, A), 1 month post *M. tb* infection (C) and 3 months post *M. tb* infection (E) showing MCs (white arrowheads) stained with anti-Acetylated tubulin (red, upper panels) and N2ICD (green, lower panels). Nuclei are stained with DAPI (blue). Yellow dotted lines indicate the basal margin of the airway epithelium. (B,D,F) Distal airway sections from control (BALB/c age matched uninfected cohort, B), 1 month post *M. tb* infection (D) and 3 months post *M. tb* infection (F) showing MCs (white arrowheads) stained with anti-Acetylated tubulin (red, upper panels) and N2ICD (green, lower panels). Nuclei are stained with DAPI (blue). Yellow dotted lines indicate the basal margin of the airway epithelium. Note that nuclear N2ICD staining is absent in MCs of distal airway in 3 months post *M. tb* infected lungs. **(G)** Volcano plot comparing gene expression in tracheal and distal airway ROIs in lungs 3 months post *M. tb* infection. Individual genes (dots) showing log2 fold change >0.5, p<0.05 (enriched in distal airway, cyan) and log2 fold change <-0.5, p<0.05 (enriched in trachea, purple) are highlighted. The number of highlighted genes in trachea and distal airway are shown in boxes in corresponding regions of the volcano plot. Top 5 enriched genes in trachea and distal airway are labelled. **(H)** Heat map showing the differential expression of genes involved in Notch signaling (MSigDB REACTOME database and KEGG database combined) in trachea and distal airway of *M. tb* infected animals compared to control. Note the decrease in expression of the Notch pathway genes and Notch2 in particular in the distal airway post infection. Genes showing a statistically significant change are marked with asterisk (*). **(I)** Graphical summary of the perturbations in ciliary lengths and the P-D gradient reported here and a model for the role of Notch signaling in MCs in stabilizing ciliary length. Notch signaling maintains a balance between ciliary elongation and retraction. The absolute length to which cilia grow is likely dependent on other signals (see Discussion).

To further examine the status of Notch signaling post *M. tb* infection, we once again adopted a spatial transcriptomic approach. Sections from trachea and lung from *M. tb* infected animals (3 months post infection) were analysed using the GeoMX DSP platform (10 ROIs from trachea, 12 ROIs from distal airway). We performed a differential gene expression analysis between distal airway and tracheal ROIs of *M. tb* infected animals. Volcano plots showed several genes that are differentially expressed across these ROIs from infected animals (genes with |log 2-fold change| > 0.5 and p < 0.05 are used) (Figure 5G). 372 genes in the tracheal ROIs and 443 genes in distal airway ROIs were significantly enriched in the respective regions of the airway (list of genes in Source Data Table 2). To probe the status of Notch signaling in *M. tb* infected lungs, we compared expression of a list of genes that are targets of the Notch signaling pathway (reference genes from MSigDB REACTOME database and KEGG database combined) in trachea and distal airway of *M. tb* infected lungs with their expression in trachea and distal airway *FoxJ1*^CreERT2/+^; Rosa^Tdtomato/+^ controls (see Figure 3) respectively (Figure 5H). We observed significant downregulation of many of these genes in distal airway 3 months post *M. tb* infection when compared with expression in distal airway of control lungs. Pertinently, levels of *Notch2* were significantly downregulated 3 months post *M. tb*. The analysis of Notch signaling post *M. tb* infection shows that active regulation of ciliary lengths indeed correlate with the downregulation of Notch signaling.

## DISCUSSION

Our efforts to probe the role of Notch signaling in MC homeostasis in the lung were stimulated by the finding that the pathway is active in all MCs throughout adult life. Our findings highlight the interplay between signaling pathways, ciliary architecture, and environmental challenges in maintaining respiratory function. We report that downregulation of Notch signaling in MCs, either by targeted inhibition of the pathway or induced by *M. tb* infection, lead to changes in ciliary length and perturb the P-D ciliary gradient. The salient implication is that ciliary lengths and the ciliary gradient in the lung are actively controlled and Notch signaling is the stabilizing mechanism. In the discussion that follows we comment on the roles of the Notch pathway in MC development and homeostasis, on the relevance of the findings reported here to the human lung, and on the functional implications of active control of ciliary length (and the P-D ciliary gradient).

The Notch pathway has multifarious roles during MC development and renewal (22, 23). Downregulation of signaling in uncommitted progenitors by miR-34/449 initiates the program of multiciliogenesis (24). A recent study suggests that the pathway has a distinct role later in MC development. At late stages, Notch signaling contributes toward establishing a P-D gradient of Prominin 1 expression in committed MCs. It is thought that Prominin 1 may inhibit ciliary growth in a dose-dependent manner to establish differences in ciliary length and set up a P-D gradient (8). Whether the above-mentioned roles of Notch signaling in MC development are relevant to the role of the pathway in MC homeostasis is not clear. Spatial transcriptomics of tracheal and distal airway post Notch inhibition (and upon *M. tb* infection) do not support the model that Notch signaling directly controls ciliary elongation or retraction via concerted transcriptional control of ciliogenic or ciliary genes (Figure 3). We have also probed the expression of Prominin 1 in the adult lung. Spatial transcriptomic analysis of Prominin 1 expression suggests that there is a gradient of mRNA expression in the airways with higher levels of expression in the distal airway (Distal Airway:Trachea Log2 FC=1.48, p=0.01). The expression of Prominin 1 in the trachea is increased in *Rbpjk*-flox/flox (*Rbpjk*-flox/flox Trachea:Control Trachea Log2 FC=0.5, p=0.01). These are consistent with the possibility that increased Prominin 1 expression reduces ciliary length. However, we also note that levels of Prominin 1 expression are unaffected in the distal airway in *Rbpjk*-flox/flox (*Rbpjk*-flox/flox distal airway: Control distal airway Log2 FC=-0.18, p=0.5). Moreover, Prominin 1 expression in trachea and distal airway are unaffected in *M. tb* infected lungs (Distal Airway:Trachea Log2 FC=0.11, p=0.6). It appears that increases in ciliary length in the distal airway, either in *Rbpjk*-flox/flox or in *M. tb*, are not associated with decreased levels of Prominin 1. Nevertheless, the role of Prominin 1 in the regulation of ciliary length during homeostasis merits a deeper investigation.

We propose that the role of Notch signaling in MCs during homeostasis is to maintain a balance between the processes of ciliary elongation and retraction and establish a steady state (summarized in Figure 5I). In our efforts to probe the role of Notch signaling in MC homeostasis, we also examined how overexpression of the Notch 1 intracellular domain (NICD) in the airways, under the control of an inducible Nkx2.1creER, impacted MCs 10 d and 2 months post induction. The overexpression of NICD under Nkx2.1creER is expected to upregulate Notch signaling in many MCs (NJ, AG, unpublished). We did not detect any obvious alteration of ciliary lengths in NICD overexpressing lungs (NJ, AG, unpublished). This supports the model that Notch signaling acts to stabilize ciliary length and that there are other signals that regulate ciliary length. Along the same lines, perturbations to Notch signaling have opposite effects on ciliary length in the proximal and distal airway. Cilia in the proximal airways shorten while cilia in the distal airway elongate. Together, these observations suggest that there are signals other than Notch that regulate ciliary length.

Histologic analyses of MCs in human lungs of smokers, of asthmatics and COPD patients, all reveal a reduction in ciliary length (15,25–27). A study examining MCs located in the nasal passages in patients with nasal polyps shows that these cells exhibit increased ciliary length (28). We propose that the spectrum of phenotypes in MCs upon Notch inhibition are relevant to the changes observed in the human lung and that the downregulation of Notch signaling could underlie many of these changes.

Studies on the regulation of ciliary beating demonstrate that the cilia in the trachea and upper airways have the capacity to tune their beat frequencies in response to chemical cues (6,7). Our study reveals yet other mechanisms by which MCs, and the airway mucociliary escalator, can respond to environmental challenges. The finding that Notch signaling in MCs stabilizes ciliary length posits that the structure and function of the airway mucociliary escalator is actively controlled via Notch signaling to keep the airways clear. It is plausible that changes in ciliary lengths and the P-D gradient will alter the rates of fluid flow and the efficiency of mucociliary clearance through the respiratory tract. Future studies will attempt to recapitulate the effects of Notch inhibition in tracheal and distal airway MCs *in vitro*. These models will facilitate both mechanistic studies on how ciliary lengths are regulated and functional studies on the possible impact of altered ciliary lengths on mucociliary clearance.

## MATERIALS AND METHODS

### Study design

The main aim of the study is to elucidate the role of Notch signaling in MCs during homeostasis and upon pathogen challenge. We inhibited Notch signaling using two approaches, first a systemic inhibition using inhibitory antibodies against ligands for Notch receptors (Jagged1 and Jagged2). Second, using a genetic approach by conditionally deleting *Rbpjk*, in FOXJ1 expressing MCs and investigating the transcriptional changes using spatial transcriptomics approach. We carefully looked at the ciliary architecture and P-D ciliary gradient in different regions of the airways. Mice model of *M. tb* infection was used to probe the impact of pathogen challenge in ciliary remodelling.

### Animal ethics and Animal handling

All the experiments involving mice were conducted either at inStem, Bangalore or FNDR (Foundation for Neglected Disease Research), Bangalore or at the National Gnotobiotic Rodent Resource Center (NGRRC) at the University of North Carolina (UNC), Chapel Hill. For experiments conducted at inStem and FNDR, the procedures were reviewed and approved in advance by the Institutional Animal Ethical Committee in accordance with guidelines established by the Committee for Control and Supervision of Experiments on Animals (CCSEA) (FNDR registration number -2082/PO/Rc/S/2019/CCSEA). For experiments conducted at UNC, the studies and protocols were approved by the Institutional Animal Care and Use Committee for the University of North Carolina in accordance with the guidelines outlined by the Animal Welfare and the National Institutes of Health.

Groups of 3-5 age matched adult animals that were 6-8 weeks of age were used for all the experiments. Female BALB/c animals were used for all *M. tb* infection experiments. For all other experiments, animals of either gender, males or females, were used depending on the available genotype. Any procedure that could conceivably cause distress to the animals employed peri-procedure anaesthesia with isoflurane gas (Baxter Healthcare Corp.) delivered by an aesthetic vaporizing machine. In addition, all animals were monitored for signs of distress and euthanized if in distress. Euthanasia was performed by anesthetizing the animals followed by cervical dislocation.

### Mouse strains

All animal strains were housed under Specific Pathogen Free environment (SPF) conditions in the respective animal facilities at the Bangalore Life Science Cluster, FNDR and NGRRC. C57BL/6J (JAX# 000664), *FoxJ1*^CreERT2^ (JAX# 27012), BALB/c and Rosa26 Ai14 td-tomato (JAX# 007914) were obtained commercially from The Jackson Laboratory. Floxed *RBP-J* strain (RIKEN# RBRC01071) was a kind gift from Dr. Mitsuru Morimoto, RIKEN. Frozen embryos of the Floxed *RBP-J* strain were cryo-rederived at the Mouse Genome and Engineering Facility at the National Centre for Biological Sciences. *FoxJ1*^CreERT2/+^ strain was crossed to Floxed *RBP-J* and Rosa26 Ai14 Td-tomato strains to generate *FoxJ1*^CreERT2/+^; *Rbpjk*^flox/flox^ and *FoxJ1*^CreERT2/+^; Rosa^Tdtomato/+^ strains respectively.

#### Antibody mediated Notch inhibition

C57BL/6 mice, *FoxJ1*^CreERT2/+^; Rosa^Tdtomato/+^ mice (≥ 6 weeks of age) were injected intraperitoneally with non-targeting IgG (anti gp120) or anti-Notch2/Notch1 or anti-Jagged1/Jagged2 antibodies diluted in PBS and sacrificed at the time points indicated in the experimental schematics. Antibody concentrations used were as follows, non-targeting IgG (anti gp120) - 40 mg/kg, anti Notch1- 10 mg/kg, anti Notch2- 30 mg/kg, anti Jagged1- 20 mg/kg and anti- Jagged2- 20 mg/kg. Animals were sacrificed at respective time points as indicated in figures.

#### Genetic ablation of *Rbpjk* in multiciliated cells

*FoxJ1*^CreERT2/+^; *Rbpjk*^flox/flox^ mice (6-8 weeks of age) were injected with Tamoxifen (Sigma, 250 mg/kg body weight) four times on alternate days before sacrificing at the time points indicated in the figures.

#### Infection with Mycobacterium *tuberculosis* (*M. tb*)

6-8 weeks old female BALB/c animals were infected with H37Rv strain of *M. tuberculosis*, (ATCC27294) in a Glas-Col Aerosol Infection Chamber (Glas-Col, Terre Haute, IN, USA), calibrated to deliver ∼ 100-200 colony-forming units of *M. tb*, in a BSL3 facility. Infected mice were housed in IVC isolators (Citizen, India) during the entire period of experimentation. Lungs of infected animals (n=3 for CFU and n=5 for histology) were collected along with the healthy/naive mice controls (n=3) on 1, 30, 90, and 180 days post infection.

#### Germ-free (GF) mice

GF mice were generated in the C57BL/6J background at the CGBID Gnotobiotic core, UNC. These mice had no detectable levels of yeast, bacteria, parasites or molds. The sterility of the isolators housing the GF mice were regularly monitored. Samples from faeces, buccal-paws-cage swabs, food and drinking water were regularly examined for bacterial contamination through Gram staining and 16S PCR on sampled faeces.

### Histology, Immunofluorescence, and Imaging

Lungs were inflated with 4% (weight/volume) Paraformaldehyde + 2% low-melting agarose in PBS and immersed in the same fixative for 12 hours at 4°C. The lobes were separated and the left lobe was used for making thick sections (200 µm). The remaining lobes were incubated in PBS at 60°C overnight for melting the agarose and subsequently processed for embedding into paraffin for histological analysis. Heat mediated antigen retrieval was performed using antigen unmasking solution from Vector labs (pH9) prior to immunostaining. pH6 antigen unmasking solution was used when staining for Notch2ICD. Immunofluorescence analysis utilized the following antisera: Rabbit anti-Notch2ICD (Abcam, 1:100), Goat anti-Uteroglobin (Merck, 1:1000), Mouse anti-Acetylated tubulin (Sigma, 1:2000), Mouse anti-Foxj1 (eBioscience, 1:250), Rabbit anti-RBPUSH (Cell signaling Technologies, 1:200), Mouse anti-RFP (Abcam, 1:300), Rabbit anti-RFP (Rockland, 1:300), Rabbit anti-Acetylated tubulin (Abcam, 1:1000). Alexa 405/488/568/647- conjugated Donkey anti-mouse/rabbit/goat (Invitrogen, 1:300). Sections were imaged either on Zeiss LSM-780 or LSM-980 laser-scanning confocal microscopes.

### Quantification of cell frequencies

Cell frequencies were calculated from immunostained paraffin sections (4-5 µm). Multi-channel images were acquired using confocal microscopes and cell frequencies were obtained by manual counting in images using Fiji (ImageJ). Nuclei were identified by presence of DAPI and cells that were positive for the respective markers were counted. For CCs and MC frequencies, CC10 and FOXJ1 immunostained cells were quantified and normalized to the length of basal lamina both in trachea and distal airway. Closed airways with diameter of ≤ 200 µm or 200 µm from the bronchoalveolar duct junctions (BADJ) of open-ended airways that transition into the alveoli were considered as distal airway. All frequencies are plotted as mean ± standard deviation from n=3 animals per condition. Student’s t-test was used for calculating the significance in the statistical analysis.

### Scanning Electron Microscopy (SEM)

The left lobe from each of the animals was sectioned (200 μm) using a compresstome. Sections were washed repeatedly in phosphate buffer at 60°C for 10 min, wash to melt the agarose and fixed with 2.5% glutaraldehyde overnight at 4°C. Fixed lung sections were dehydrated with increasing concentrations of acetone and dried by critical point drying using Leica EM CPD 300. Sections were then sputter coated with gold particles using PELCO SC - 7. Electron micrographs of the lung sections were taken using the ZEISS MERLIN Compact at a magnification of 3-10 kX and an accelerator potential of 2- 4 kV.

### Measurement of cilia length

Linear measurements of individual cilia of MCs were measured manually from SEM images (30). Cilia whose entire length from base to the tip was clearly discernible were traced using the free hand option in Fiji (ImageJ) and their lengths were measured and tabulated. 200-1000 cilia per animal per condition were measured across multiple SEM images (n≥3 animals per condition). Graphs for ciliary lengths were generated in Microsoft Excel by compiling the results to calculate mean and mean± standard deviation. Student’s t-test was used for calculating the p value and significance. Closed airways with diameter of ≤100µm or 100µm from the BADJ of open-ended airways that transition into alveoli were considered for imaging cilia.

### Spatial Transcriptomics

Formaldehyde fixed, paraffin embedded (FFPE) sections of trachea and lungs (5µ) from control, *FoxJ1*^CreERT2/+^; *Rbpjk*^flox/flox^ and *M. tb* infected mice were used for spatial transcriptomic profiling. Mouse anti-Acetylated tubulin (Sigma) antibody was conjugated to CF568 using Mix-n-Stain^™^ CF^™^ 568 Antibody Labelling Kit (50-100μg), Sigma. Double positivity to Acetylated tubulin and SYTO 13 (nuclear marker) was used to identify MC containing airway epithelium and define regions of interest (ROIs). GeoMx Digital Spatial Profiler (DSP) slide preparation was performed as per manufacturer’s instructions. Slides were then incubated overnight at 37°C with Mouse Whole Transcriptome Atlas (WTA) probes (NanoString) as per the NanoString GeoMx RNA-NGS manual instructions. Slides were then washed, stained with Acetylated-tubulin-CF568 and SYTO 13 for 1 hour at room temperature, imaged on the GeoMx DSP at 20X magnification and ROIs were selected in the trachea and the distal airway. Indexed oligonucleotides from each ROI were released and deposited into a 96-well plate on exposure to 385 nm light (UV), with a microcapillary. The collected DNA oligos from each ROI were subjected to Illumina library preparation for sequencing. A total of 74 ROIs and 2 NTC were profiled. Sequencing libraries were prepared according to the NanoString GeoMx-NGS Readout Library Prep manual. The PCR reaction was performed using the NanoString SeqCode primer for 21 cycles. Libraries were sequenced on an Illumina NovaSeq 6000, with 2 x 151 bp paired end reads.

### Nanostring GeoMx WTA data analysis

The sequenced fastq files were processed further to convert into.dcc (Digital Count Conversion) format using the GeoMx NGS Pipeline v.2.3.3.10 from NanoString. The trimming of adapters, removal of duplicates and mapping to the reference probe barcodes present in the WTA panel and quantification would be performed during the conversion process of fastq to dcc files. The individual ROIs would hence result in a single dcc file. The resulting dcc files from each AOI were further imported and analyzed using the GeoMxTools (1) R package. R version 4.2 (https://www.r-project.org/about.html) was used throughout the analysis unless stated otherwise. The analysis within GeoMxTools pipeline includes QC for both Probe and Segment, Normalization, Unsupervised clustering. The parameters used during the QC of segments includes: minSegmentReads = 1000, percentTrimmed = 80, percentStitched = 80, percentAligned = 80, percentSaturation = 50, minNegativeCount = 10, maxNTCCount = 1000, minNuclei = 200, minArea=5000 and minLOQ = 2. The parameters used for the QC or filtering of probes (target genes) includes: geometric mean of each probe’s counts from all segments divided by the geometric mean of all probe counts representing the target from all segments < 0.1 and percentage of segments within which the probe is identified as an outlier based on Grubb’s test >= 20%. A probe is removed locally (from a given segment) if the probe is an outlier according to the Grubb’s test in that segment. After appropriate QC on segments and target genes, the counts were normalized using the Q3 method of normalization. The differential gene expression testing was performed using the LMM model within GeoMxTools R package, which adjusts for the fact that the multiple regions of interest placed per tissue section are not independent observations as understood by other statistical tests.

### Statistical analysis

A two**-**tailed unpaired Student’s t-test was employed to determine the statistical significance between conditions in the measurements of ciliary length and frequencies. P value ≤0.05 is considered statistically significant and is denoted by ‘*’ in the corresponding plot. All experiments were conducted independently on at least 3 animals (n≥3) and error bars indicate mean ± standard deviation. Matplotlib (python 3.0 package) was used to plot dot plots and frequency distribution.

## ACKNOWLEDGEMENTS

The team expresses gratitude to TheraCUES innovations private limited, Bangalore, for spatial transcriptomics experiment and Manju Moorthy for post sequencing analysis. Electron Microscopy Facility, Central Imaging and Flow Cytometry Facility (CIFF) and the Animal Care and Resource Centre (ACRC) at the Bangalore Life Science Cluster (BLiSC) have provided their support in conducting experiments and maintaining experimental animals. AG would like to thank Dr. Balasubramanian V for inspiration, Dr. Madan. Rao for discussions and Dr. Narmada Khare for critical reading of the manuscript.

## Funding

Institutional core funds from inStem, Bangalore.

Funding from Department of Biotechnology (DBT), Department of Science and Technology – Science and Engineering Research Board, India.

Chan Zuckerberg Initiative (Mapping the Paediatric Inhalation Interface: Nose, Mouth and Airways) (J.S.H., R.C.B., K.M.B., and A.G.).

Cystic Fibrosis Foundation Research Development Project BOUCHE19R0 (R.C.B)

National Institute of Health grants NHLBI P01HL164320 and NIDDK P30 DK065988 (R.C.B)

National Institute of Health grants NHLBI R01 HL 150541-01 (A. L-B).

Council of Scientific and Industrial Research (CSIR) 09/860(1273)/2019-EMR-I (N.J) Fellowship from DBT (S.M.L. and A.D.)

## Author contributions

Conceptualization and methodology: A.G, N.J, R.K.S, K.M.B

Performing experiments and data analysis: N.J, A.D, S.M.L, V.M.P, N.C.N, B.K, S.B, L.B.A, Z.A.M

Data curation and visualization: N.J, V.M.P, A.D, S.M.L

Resources: A.G, J.S.H, R.C.B, D.L, K.M.B, R.K.S, L.B.A

Supervision: A.G, R.K.S, S.N, K.M.B, L.B.A

Writing and correcting manuscript: A.G, N.J, S.M.L, A.D, V.M.P, L.B.A, K.M.B

## Data and materials availability

All data required to evaluate the conclusions are available in the main text or the supplementary materials.

## SUPPLEMENTARY FIGURES

**Figure S1-1:**
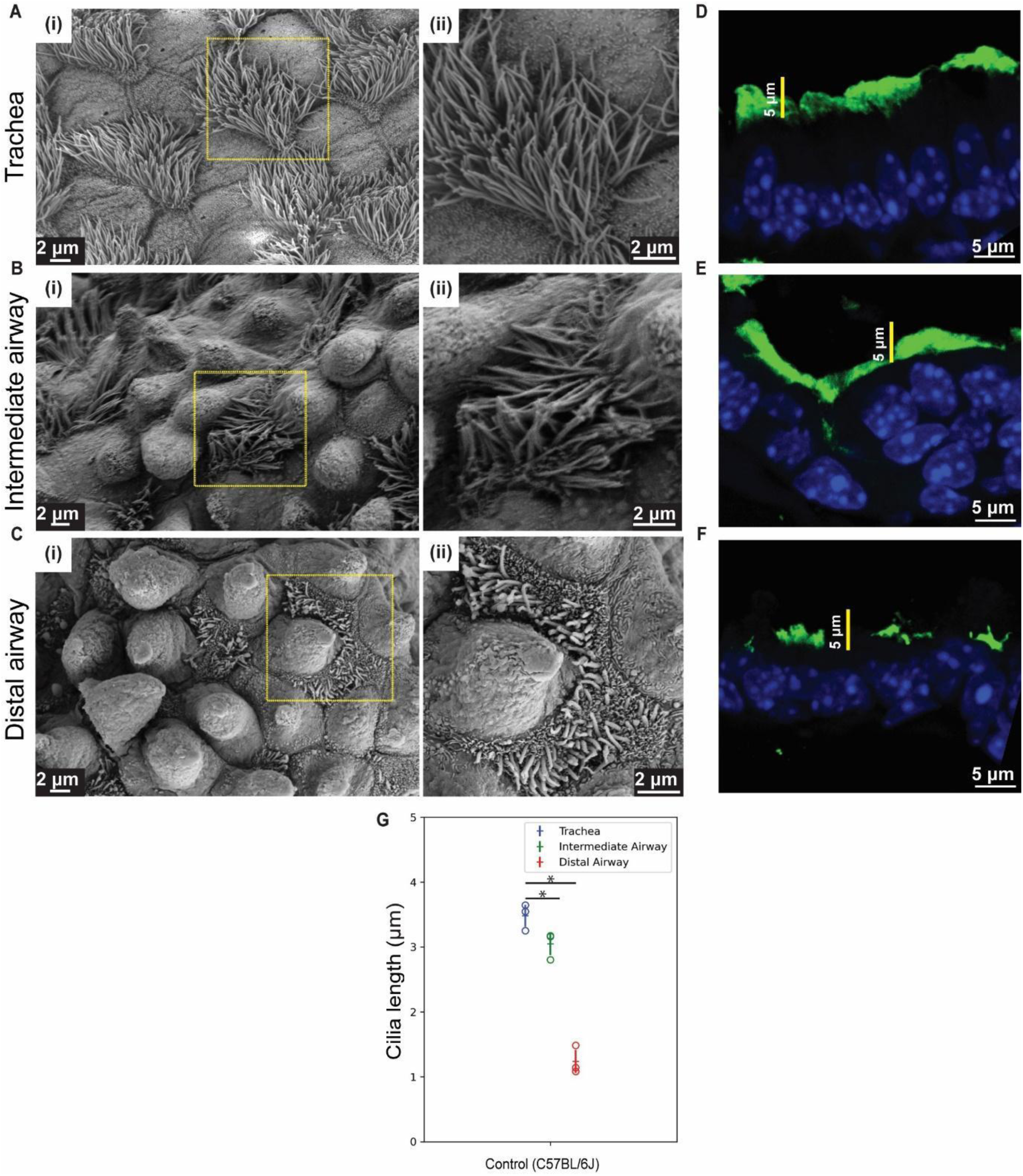
Ciliary lengths along the proximal-distal axis of airway epithelium. **(A-C)** Scanning electron micrographs showing pattern of ciliary length in MCs along the P-D airway axis. Micrographs showing MCs in trachea (A), intermediate airway (B) and distal airway (C) of 2 months old control (C57BL/6J) airway epithelium. Magnified images of boxed regions in (i) are shown in (ii). **(D-F)** Distribution of anti-Acetylated tubulin (green) in airways from control lungs. Ciliary staining in trachea (D), intermediate airways (E) and distal airway (F) of 2 months old control (C57BL/6J). Note the pattern of ciliary staining in trachea, intermediate airway and distal airway. **(G)** Quantification of ciliary length in micrographs (µm, see methods). Dot plots of ciliary lengths in trachea (blue) and distal airway (red) from 2 months old control (C57BL/6J, see text for details, n=667 cilia in trachea, n=223 cilia in intermediate airway and n=1795 cilia in distal airway). Hollow circles indicate mean ciliary length in individual animals. Data in dot plot represent mean ± standard deviation (n=3 animals), (* denotes p<0.05, Student’s t-test). Note the pattern in ciliary lengths across the P-D axis of the airway.

**Figure S1-2:**
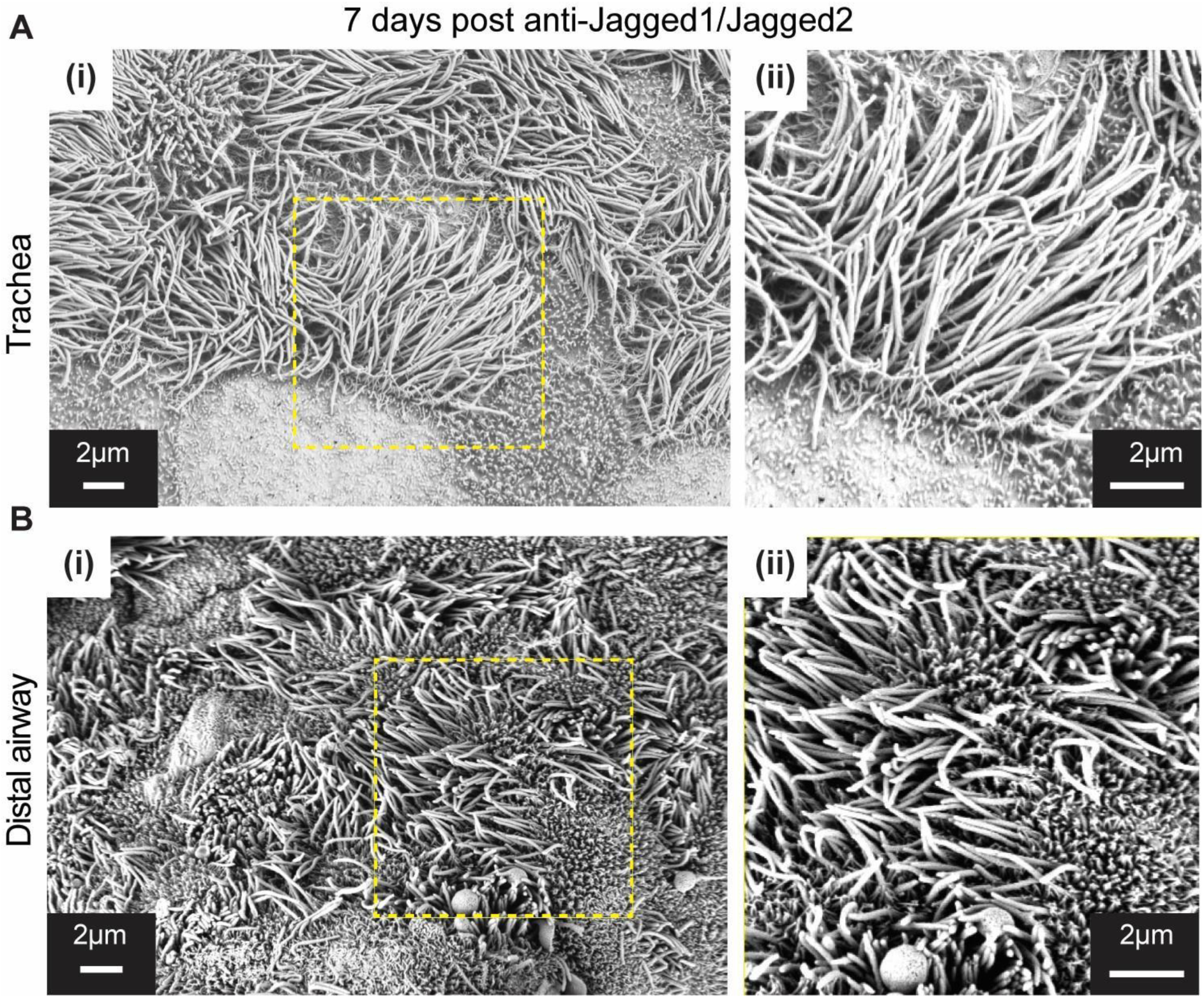
Multiciliated cell remodelling in 7 days post anti-Jagged1/ Jagged2 treatment. **(A-B)** Scanning electron microscopy of sections from 7 days post anti-Jagged1/Jagged2 treated lungs. Micrographs of tracheal (A(i)-A(ii)) and distal airway (B(i)-B(ii)) from 7 days post anti-Jagged1/Jagged2 treated lungs. Magnified images of boxed regions in (i) are shown in (ii). Note the decrease in length of cilia in trachea and an increase in length in distal airway by 7 days post anti-Jagged1/Jagged2 treated animals compared to control (see Figure 1 for control micrographs).

**Figure S1-3:**
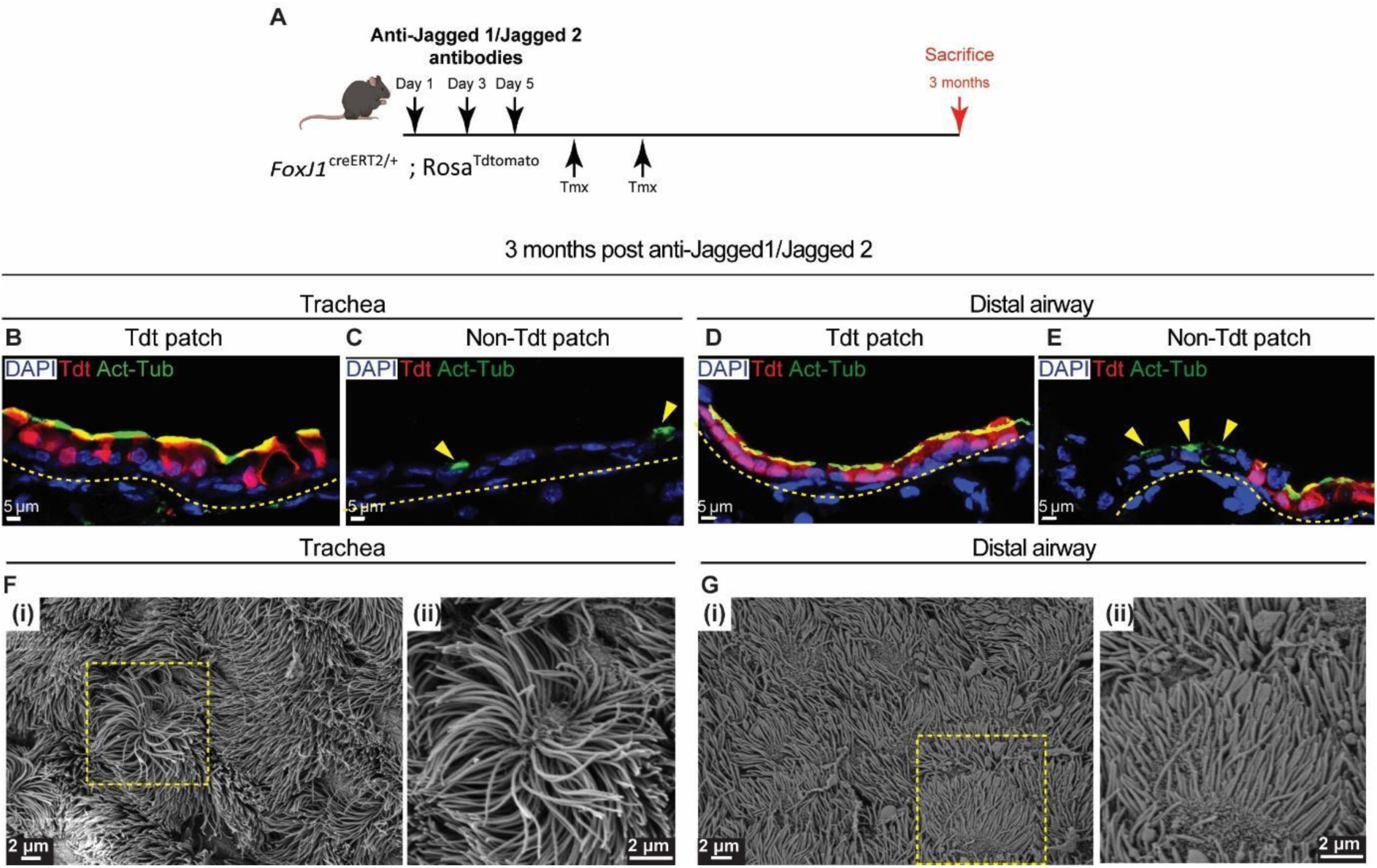
Fate of MCs and motile cilia 3 months post anti-Jagged1/Jagged2 treatment. **(A)** Diagram showing experimental pipeline for anti-Jagged1/Jagged2 antibody treatment in *FoxJ1*^CreERT2/+^; Rosa^Tdtomato/+^ mice. **(B-C)** Immunofluorescence image showing MCs distribution in trachea, 3 months post anti-Jagged1/Jagged2 treatment. (B) Immunofluorescence image showing MCs in trachea stained with Tdt (red, ‘old MCs’ co-stained with anti-Acetylated tubulin, green) identified by lineage tracing (see text for details). (C) Immunofluorescence image showing MCs in trachea which are Tdt negative suggesting they are newly formed MCs. **(D-E)** Immunofluorescence image showing MCs distribution in distal airway, 3 months post anti-Jagged1/Jagged2 treatment. (D) Immunofluorescence image showing MCs in distal airway stained with Tdt (red, ‘old MCs’ co-stained with anti-Acetylated tubulin, green) identified by lineage tracing. (E) Immunofluorescence image showing MCs in distal airway which are Tdt negative suggesting they are newly formed MCs. Yellow dotted lines indicate the margin of airway epithelium. **(F-G)** SEM images showing old MCs of 3 months post anti-Jagged1/Jagged2 treated lungs. Micrographs of trachea (F(i)-F(ii)) and distal airway (G(i)-G(ii)) of 3 months post anti-Jagged1/Jagged 2 treated animals. Magnified images of boxed regions in (i) are shown in (ii). Note that the representative SEM images are sourced from regions which only have MCs and no CCs, which represents the old MC population (Tdt positive MCs, see text for details).

**Figure S2-1:**
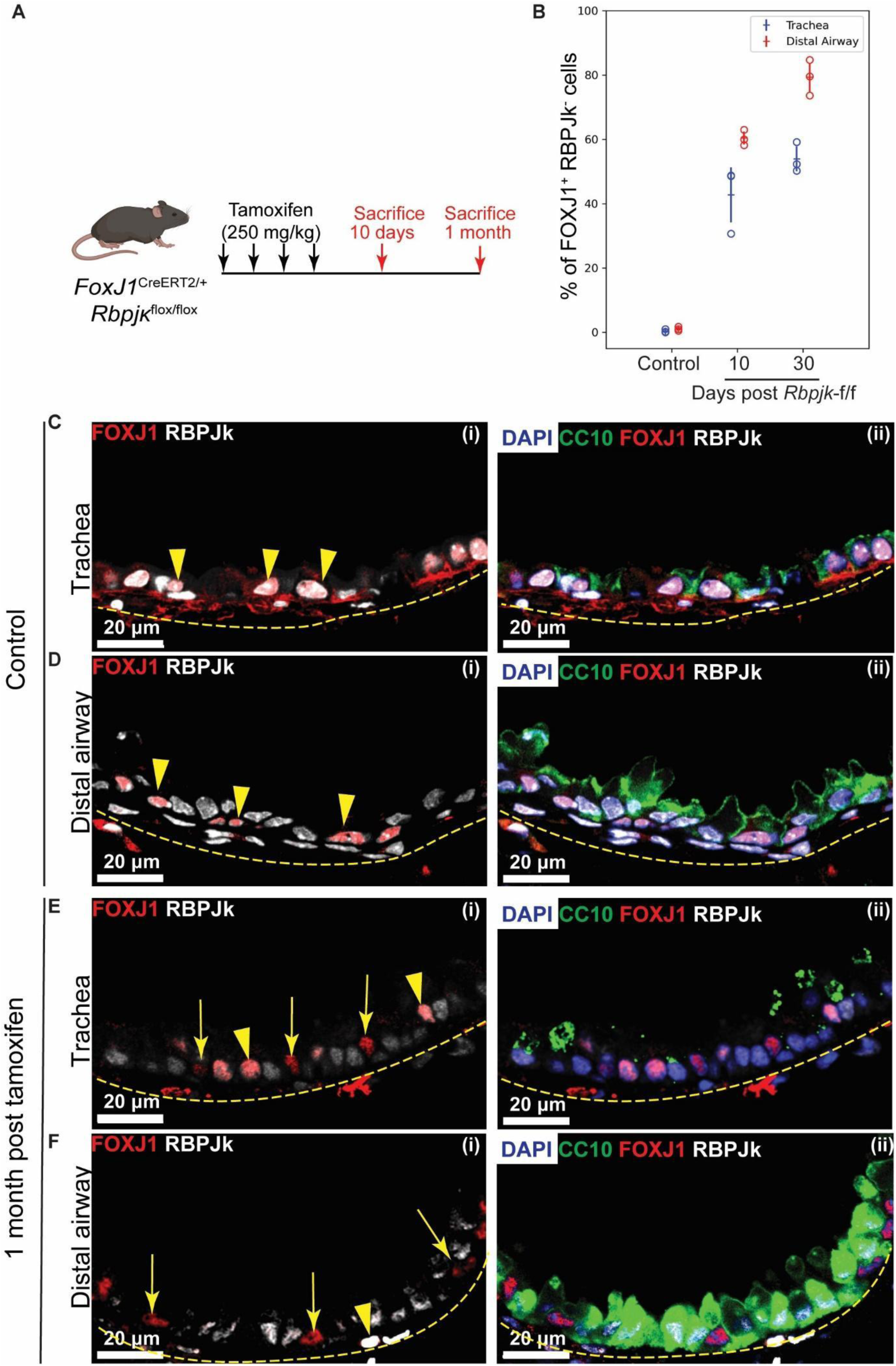
Efficiency of genetic ablation of RBPJκ in MCs. **(A)** Genetic strategy and tamoxifen regimen to conditionally deplete *Rbpjκ* in MCs. **(B)** Dot plot showing frequencies of FOXJ1+ RBPJκ-cells in trachea (blue) and distal airway (red) of control (age matched uninduced littermate), 10 days and 1 month post tamoxifen induced (“*Rbpjk*-flox/flox”) lungs. Hollow circles indicate mean FOXJ1+ RBPJk- cell frequency in immunostained sections from an individual animal (see text, methods). Data in the dot plot represent mean ± standard deviation (n= 3 animals). **(C-F)** Immunofluorescence images showing the efficiency of *Rbpjk* ablation in MCs. Immunostaining for MC specific marker FOXJ1 (red), RBPJk (grey), CC marker CC10 (green) and nuclear stain DAPI (blue) in control (age matched uninduced littermate) and 1 month post tamoxifen induced (*Rbpjk*-flox/flox) lungs. RBPJk (grey) stainings in MCCs (FOXJ1, red) of control trachea (C(i)-C(ii)) and distal airway (D(i)-D(ii)) and 1 month post tamoxifen, trachea (E(i)-E(ii)) and distal airway (F(i)-F(ii)). Yellow arrowheads point towards MCs which are double positive for FOXJ1 and RBPJk and yellow arrows point towards MCs that lack RBPJk. Yellow dotted lines indicate the margin of airway epithelium. Note that the majority of the MCs are devoid of RBPJk in distal airway 1 month post tamoxifen induction.

**Figure S2-2:**
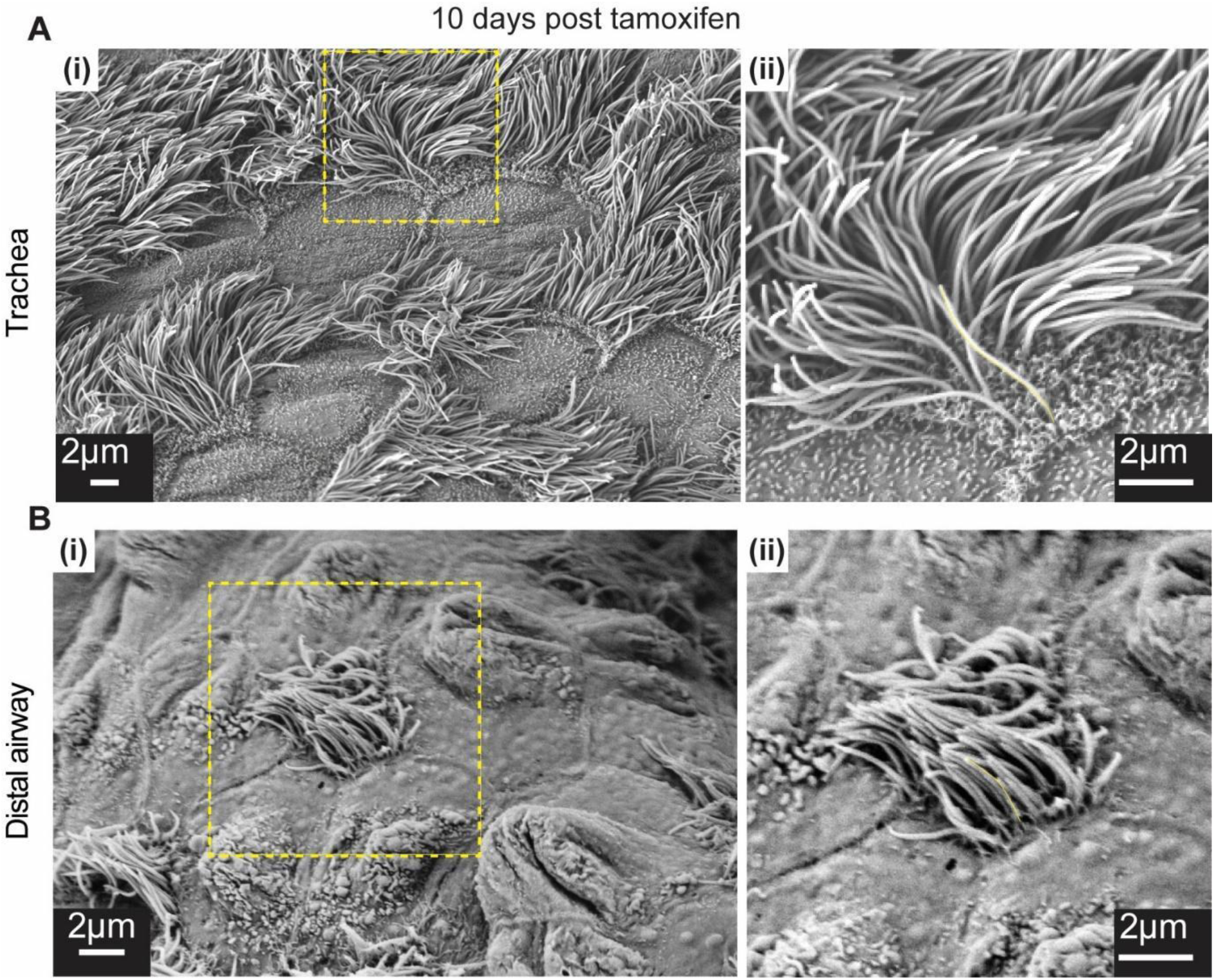
Multiciliated cell remodelling in 10 days post tamoxifen treated *Rbpjk*-flox/flox animals. **(A-B)** Scanning electron microscopy of 10 days post tamoxifen treated airway epithelium. Micrographs of tracheal (A(i)-A(ii)) and distal airway (B(i)-B(ii)) from 10 days post tamoxifen treated lungs. Magnified images of boxed regions in (i) are shown in (ii). Note the increase in length in distal airway by 10 days post tamoxifen treatment compared to control. (see Figure 2 for control micrographs).

**Figure S4-1:**
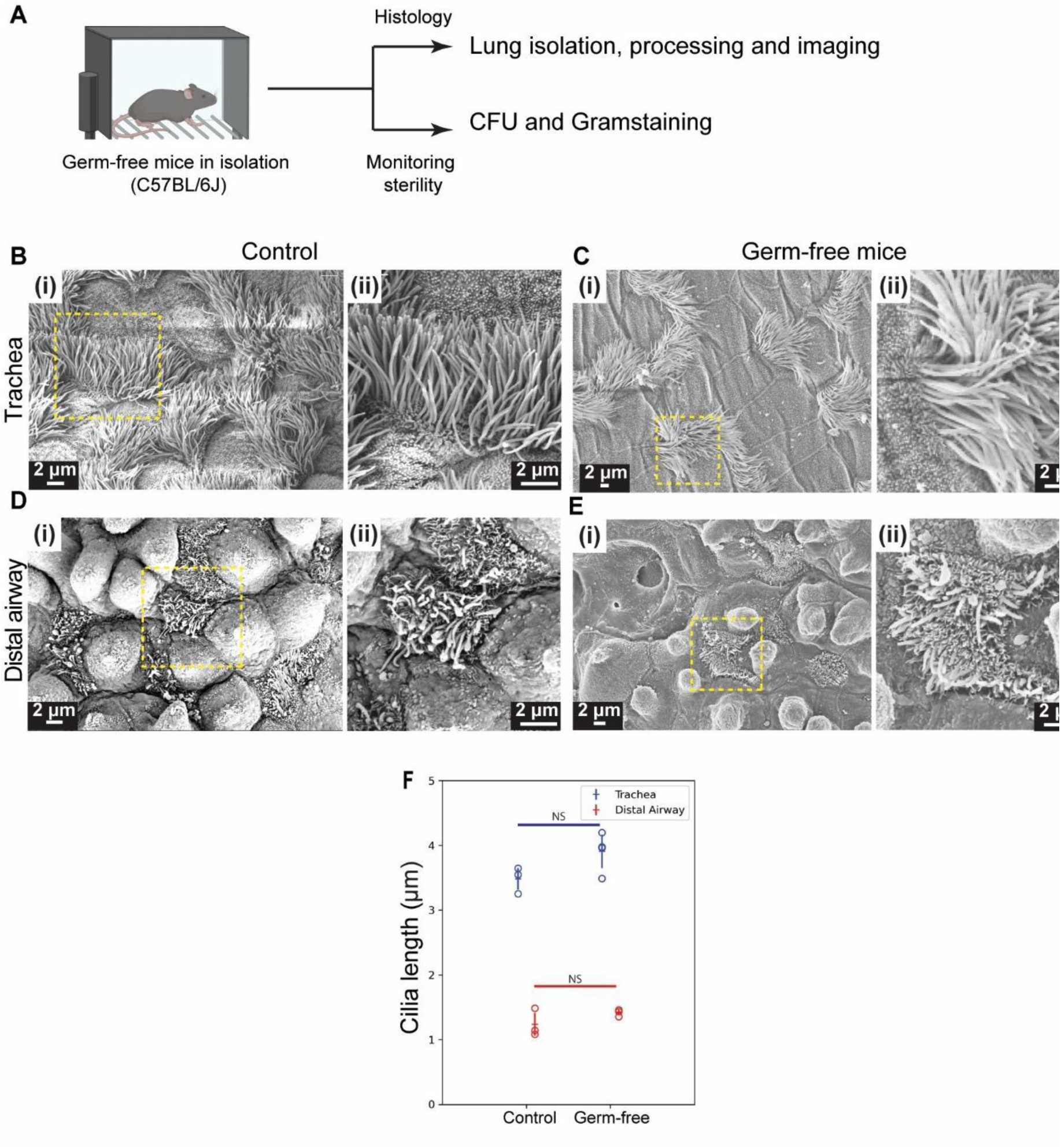
Phenotype of multiciliated cells is unaffected in lungs from germ-free mice. **(A)** Blueprint for generating germ-free mice. **(B-E)** Scanning electron micrographs showing cilia of MCs in trachea and distal airway of germ-free mice and age matched control (C57BL/6J). **(B,D)** Micrographs of trachea (B(i)-B(ii)) and distal airway (D(i)-D(ii)) from age matched control (2 months old). Magnified images of boxed regions in (i) are shown in (ii). **(C,E)** Micrographs of trachea (C(i)-C(ii)) and distal airway (E(i)-E(ii)) from germ-free mice. Magnified images of boxed regions in (i) are shown in (ii). **(F)** Dot plots of ciliary length in trachea (blue) and distal airway (red) from age matched control lungs (n=667 cilia in trachea and n=1795 cilia in distal airway) and germ-free lungs (n=1022 cilia in trachea and n=1883 cilia in distal airway). Hollow circles in the dot plot indicate mean ciliary length of individual animals. Data in dot plot represent mean ± standard deviation (n= 3-4 animals), (*denotes p<0.05, Student’s t-test).

**Figure S4-2:**
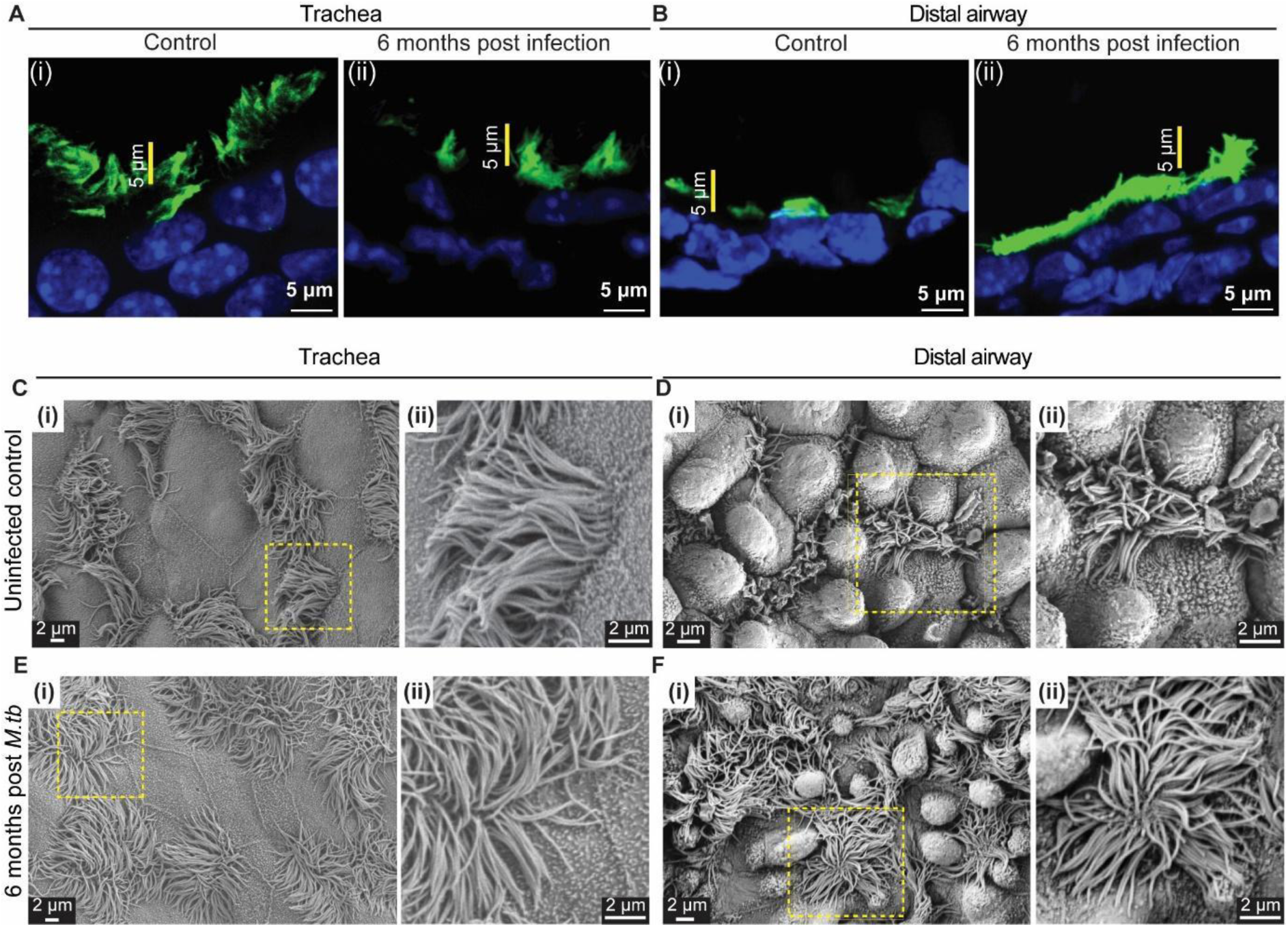
Ciliary morphology 6 months post *M. tb* infection. **(A-B)** Ciliary staining in uninfected age matched control and 6 months post *M. tb* infected lung. (A) Uninfected control trachea (A(i)) and 6 months post *M. tb* infected trachea (A(ii)) stained with anti-Acetylated tubulin (green). (B) Uninfected control distal airway (B(i)) and 6 months post *M. tb* infected distal airway (B(ii)) stained with anti-Acetylated tubulin (green). Note the increase in height of ciliary band in 6 months post *M. tb* infected distal airway compared to control. **(C-F)** Scanning electron microscopy of sections from uninfected control and *M. tb* infected lungs. **(C-D)** Micrographs of trachea (C(i)-C(ii)) and distal airway (D(i)-D(ii)) from age matched control lungs. Magnified images of boxed regions in (i) are shown in (ii). **(E-F)** Micrographs of trachea (E(i)-E(ii)) and distal airway (F(i)-F(ii)) from lungs 6 months post *M. tb* infection. Magnified images of boxed regions in (i) are shown in (ii).

